# Cortical granularity shapes the organization of afferent paths to the amygdala and its striatal targets in nonhuman primate

**DOI:** 10.1101/2021.05.06.442678

**Authors:** AC McHale, YT Cho, JL Fudge

**Author notes:** Correspondence to: Julie L. Fudge, M.D. Departments of Neuroscience, and Psychiatry The Del Monte Institute for Neuroscience University of Rochester Medical Center.

## Abstract

The prefrontal cortex (PFC) and insula, amygdala, and striatum form interconnected networks that drive motivated behaviors. We previously found a connectional trend in which granularity of the ventromedial and orbital PFC/insula predicted connections to the amygdala, and also the breadth of amygdalo-striatal efferents, including projections beyond the ’classic’ ventral striatum. To further interrogate connectional relationships among the cortex, amygdala, and striatum, and to further define the ’limbic (amygdala-recipient) striatum’, we conducted tract tracing studies in two cohorts of *Macaques* (Male n = 14, Female n = 1). We focused on the cortico-amygdalo-striatal (indirect) and cortico-‘limbic’ striatal (direct) paths originating in the entire PFC and insula. Larger data sets and a quantitative approach revealed ’cortical rules’ in which cortical granularity predicts the complexity and location of projections to *both* the basal nucleus of the amygdala and striatum. Remarkably, projections from ’cortical-like’ basal nucleus to the striatum followed similar patterns. In both ’direct’ and ’indirect’ paths to the ’limbic’ striatum, agranular cortices formed a ’foundational’, broad projection, and were joined by inputs from progressively more differentiated cortices. In amygdalo-striatal paths, the ventral basal nucleus was the ‘foundational’ input, with progressively more dorsal basal nucleus regions gradually adding inputs as the ’limbic striatum’ extended caudally. Together, the ‘indirect’ and ‘direct’ paths followed consistent principles in which cortical granularity dictated the strength and complexity of projections at their targets. Cluster analyses independently confirmed these connectional trends, and also highlighted connectional features that predicted termination in specific subregions of the basal nucleus and ’limbic’ striatum.

**Significance Statement:** The ’limbic system’ broadly refers to brain circuits that coordinate emotional responses. Here, we investigate circuits of the amygdala, which are involved in coding the emotional value of external cues, and their influence on the striatum. Regions of prefrontal cortex and insula form gradients of overlapping inputs to the amygdala’s basal nucleus, which feed forward to the striatum. Direct cortical inputs to these ’amygdala-recipient’ striatal areas are surprisingly organized according to similar principles, but subtly shift from the classic ventral striatum to the caudal ventral striatum. Together, these distinct subsystems—cortico-amygdala-striatal circuits and direct cortico-striatal circuits— provide substantial opportunity for different levels of internal, sensory, and external experiences to be integrated within the striatum, a major motor-behavioral interface.

## Introduction

The prefrontal cortex (PFC) and insula, amygdala, and striatum form interconnected networks that drive motivated behaviors (Van Hoesen et al., 1981; Yeterian et al., 2012). Specific cortical regions project to the amygdala, modulating amygdala neural responses (Klavir et al., 2013; Likhtik et al., 2014). At the same time, these cortical regions and the amygdala form uni-directional inputs to the striatum to mediate goal-directed responses (Haber and Knutson, 2010). This ‘triadic’ circuitry is implicated in human neuropsychiatric diseases (Akil et al., 2018; Ressler, 2020). Yet, organizational principles of this classic ‘limbic’ triad are still not completely understood at the cellular level in the nonhuman primate brain.

Although they are separate lobes of cortex, the PFC and insula share a progressive change in cortical laminar differentiation (Mesulam and Mufson, 1982; Barbas and Pandya, 1989). The degree of ‘granularity’ across the cortical mantle generally refers to the relative presence or absence of ‘granular cell’ layer IV (Brodmann, 1909; Carmichael and Price, 1994; Petrides and Pandya, 1999, 2002). “Agranular” cortex lacks a granular layer IV, “dysgranular” cortex has an incompletely developed layer IV, and “granular” cortex has a well-differentiated layer IV. Within these broad categories, incremental changes in laminar organization exist, leading to gradual shifts from agranular to dysgranular to granular across the cortical mantle. Classic subdivisions of the PFC and the insula are therefore not independent regions, but rather reflect this continuous laminar organization (Barbas and Garcia-Cabezas, 2016).

The basal nucleus of the amygdala is a key recipient of top-down inputs from the PFC and insula, and is a major output to the striatum (Carmichael and Price, 1995; Ghashghaei and Barbas, 2002; Cho et al., 2013). This nucleus is expanded in primates (Stephan et al., 1987), and has an increased cytoarchitectural complexity along its dorsal-ventral axis. In monkey and human, three large subdivisions of the basal nucleus are recognized: the magnocellular (Bmc), intermediate (Bi,) and parvicellular (Bpc). As their names imply, these dorsally-to-ventrally arranged basal nucleus subdivisions are distinguished by pyramidal cell size and packing density (Braak and Braak, 1983).

Amygdala inputs, particularly from the basal nucleus, are a defining input to the ’classic’ ventral striatum, which is involved in forming goal-directed behaviors (Robbins et al., 1989; Popescu et al., 2009). Prior work however has demonstrated basal nucleus projections extend past the ’classic’ ventral striatum into caudal aspects of the striatum, which we term the ‘caudal ventral striatum’ (Russchen et al., 1985; Fudge et al., 2002; Cho et al., 2013).

In a previous study examining ‘cortico-amygdalo-striatal’ paths limited to the ventromedial and orbital PFC and insula, we found evidence that relatively higher levels of laminar organization in cortico-amygdala projections governed the topography of cortico-amygdala paths, and was also related to the extent of projections in the next limb of the circuit, the amygdalo-striatal path (Cho et al., 2013). Findings from this study raised subsequent questions around whether the most differentiated dorso-lateral regions of the PFC and insula followed this topography of inputs, and how the topography of cortico-amygdala-striatal circuits compared to the topography of direct cortico-striatal circuits in striatal regions receiving amygdala input.

Here, we took a systems-based approach to quantitatively compare the organization of ’indirect’ cortico-amygdala-striatal circuits and ’direct’ cortico-striatal circuits in striatal regions receiving amygdala inputs. We first examined the entire extent of the ’indirect’ path, using multiple small injection sites of bi-directional tracer focused on the specific basal nucleus subregions. In the second set of experiments, we placed small injections of retrograde tracer into territories of ‘limbic’ (amygdala-recipient) striatum defined by the first cohort, and compared whether direct cortical-‘limbic’ striatal paths followed similar connectivity principles as indirect paths. Remarkably, we found that a recursive set of hierarchically organized connections, governed by cortical granularity, dictate both the direct and indirect cortical paths to the ‘limbic’ striatum. This organization resulted in unique connectional ’fingerprints’ in subregions of the basal nucleus of the amygdala and the ’limbic’ (amygdala-recipient) striatum, suggesting functional differences.

## Materials and Methods

### Study design

Two cohorts of *Macaca fascicularis* (n = 15 total; Male n = 14, Female n = 1, average weight: 4.46 kg, weight range = 3.3 kg - 9.3 kg, average age: 4.51 - 4.64 years, age range = 3 years – 7 years) were used for this study (Labs of Virginia, Three Springs Laboratories, and Worldwide Primates) (Table 1). In the first cohort (Cohort 1, n = 8 animals, all male; see Table 1), a bidirectional tracer injection was placed in various subdivisions of the basal nucleus of the amygdala (Fig. 1). Cohort 1 data includes new analyses in animals used in a previous study (Cho et al., 2013), as well as new cases. To examine the cortico-amygdalo-striatal circuit, we charted and analyzed the number of retrogradely labeled cells in the entire PFC/insula, and the distribution of anterogradely labeled fiber terminals in the striatum. In the second cohort (Cohort 2, n = 8 animals, 1 female, 7 males; Table 1), we used anterograde tracing maps from Cohort 1 to guide placement of retrograde tracer injections into the ‘limbic’ striatum (defined as striatal regions receiving amygdala input). Retrogradely labeled cells in the PFC/insula and in the amygdala were then charted, quantified, and analyzed.

**Figure 1:**
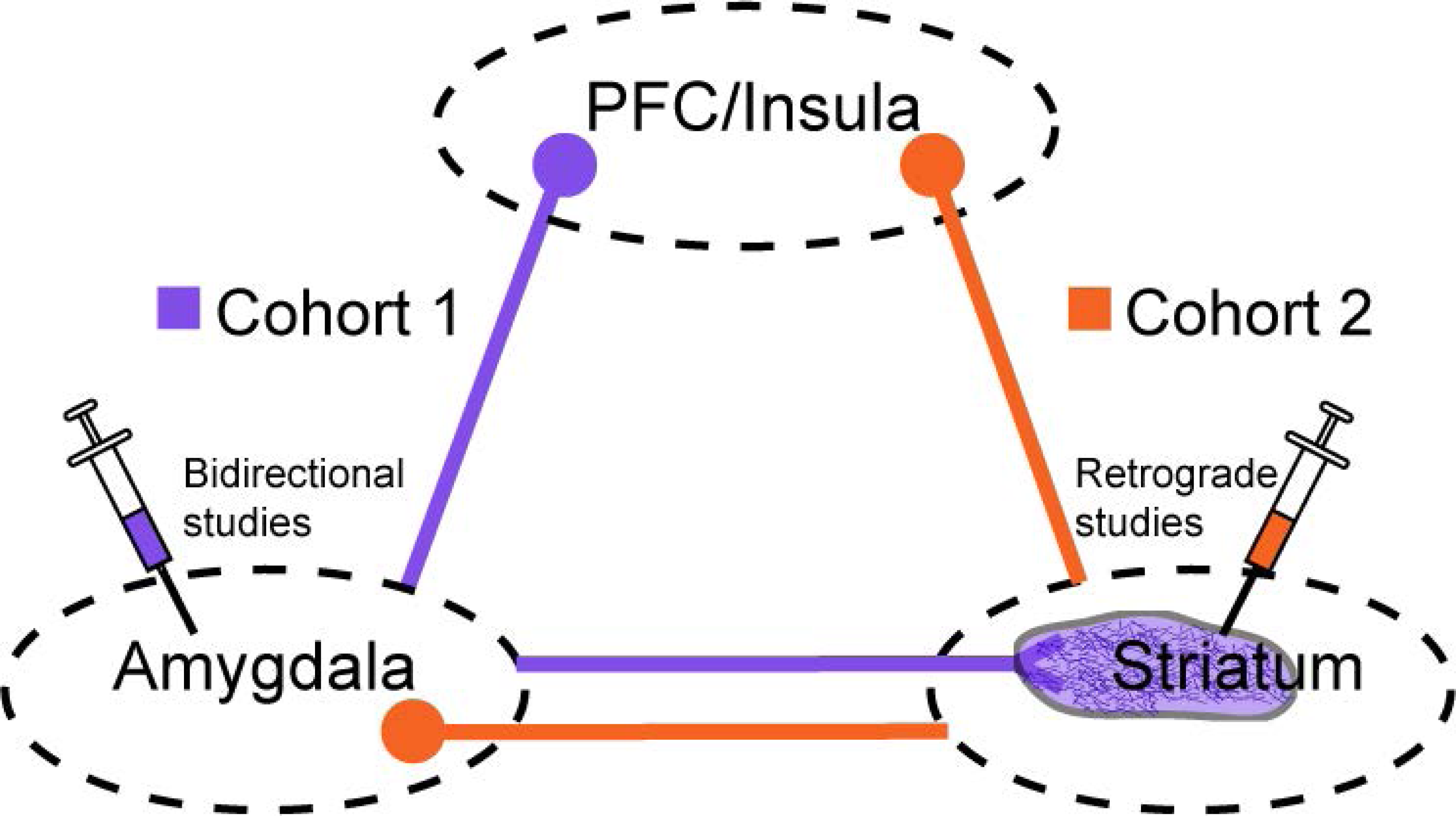
Study design. Cohort 1 animals (bi-directional tracer studies, purple) had a series of injections placed in different subdivisions of the basal nucleus of the amygdala. Resulting retrogradely labeled cells in the PFC and Insula were quantified; anterogradely labeled fibers in the striatum were mapped to guide injections in Cohort 2. Cohort 2 animals (retrograde studies, orange) had a series of injections placed in different regions of the striatum that were ‘amygdala-recipient’ (i.e. striatal regions with labeled fibers in Cohort 1 animals). Retrogradely labeled cells in the amygdala, and the PFC and Insula, were quantified.

**Table 1:** Injection site summary and monkey characteristics

| Table 1: Injection site summary and monkey characteristics |  |  |  |  |  |
| --- | --- | --- | --- | --- | --- |
| Monkey Identification | Number of injections | Injection type | Cohort Affiliation | Weight (kg) | Age (years) |
| J14 | 2 | FR<br>FS | Cohort 1<br>Cohort 1 | 3.3 | 4 |
| J15 | 2 | LY<br>FS | Cohort 1<br>Cohort 1 | 3.3 | 3-4* |
| J16 | 1 | LY | Cohort 1 | 3.6 | 3-4* |
| J20 | 1 | LY | Cohort 1 | 9.3 | 7 |
| J44 | 1 | FR | Cohort 1 | 4.4 | 4.2 |
| J47 | 1 | FS | Cohort 1 | 4.6 | 5.3 |
| J52 | 1 | LY | Cohort 1 | 4.1 | 3.5 |
| J12 | 3 | LY<br>FR<br>WGA | Cohort 1<br>Cohort 1<br>Cohort 2 | 6 | 6.6 |
| J8+ | 1 | WGA | Cohort 2 | 3.5 | 4.2 |
| J11 | 1 | WGA | Cohort 2 | 4.1 | 4.5 |
| J13 | 1 | WGA | Cohort 2 | 3.8 | 3.9 |
| J24 | 1 | WGA | Cohort 2 | 3.9 | 4.9 |
| J35** | 1 | FR | Cohort 2 | 4.5 | 3.6 |
| J41 | 1 | WGA | Cohort 2 | 4.1 | 5.8 |
| J42** | 1 | WGA | Cohort 2 | 4.4 | 4.1 |

"+" female monkey
"\*" approximate age
"\*\*" control injections

### Surgical procedures and tissue preparation

All surgeries were approved by the University of Rochester Committee on Animal Research and follow the National Institutes of Health guidelines. To determine surgical coordinates, some animals had an individualized MRI T2 scan prior to surgery; others had intra-operative electrophysiologic mapping (Fudge et al., 2004; Fudge et al., 2005; Cho et al., 2013; Decampo and Fudge, 2013).

Monkeys that underwent individualized MRI T2 scans were first administered an intramuscular injection of ketamine (10 mg/kg), then intubated with isofluorane gas during the scanning process. Three days prior to stereotactic surgery, the animals received daily prophylactic pain control with gabapentin (oral dose), which was maintained for 3 days post-operatively. On the day of surgery, monkeys were administered an intramuscular injection of ketamine (10 mg/kg), intubated, and then administered either intravenous pentobarbital (initial dose, 20 mg/kg, i.v.) (for animals undergoing electrophysiologic mapping) or isofluorane gas anesthesia. Once monkeys were placed in a stereotactic head frame, a craniotomy was performed under sterile conditions. For some animals, electrophysiological mapping was conducted first, and electrophysiological features of cells were noted during several penetrations. Injection sites were then plotted for subsequent tracer injections.

For Cohort 1, the basal nucleus was pressure-injected at various dorsoventral and rostrocaudal sites with either 40 nl of the bidirectional tracers Lucifer yellow conjugated to dextran amine (LY; Invitrogen), Fluorescein conjugated to dextran amine (FS; Invitrogen), or Fluoro-Ruby (tetramethylrhodamine) conjugated to dextran amine (FR; Invitrogen) (Table 1). Antibodies to these tracers do not cross-react, and have similar retrograde and anterograde properties, making them suitable for simultaneous use within the same animal, based on our previous work (Haber et al., 2000; Fudge et al., 2002). For Cohort 2, one striatal site per animal was pressure-injected with 40 nl of wheat germ agglutinin conjugated to horseradish peroxidase (WGA-HRP; Sigma) (Table 1). Following all surgeries, the bone flap was replaced and the overlying tissue sutured. For both cohorts, animals were deeply anesthetized and sacrificed via intracardiac perfusion 10 to 12 days after surgery [0.9% saline solution containing 0.5 ml of heparin sulfate followed by cold 4% paraformaldehyde in 0.1M phosphate buffer (PB (0.1M PO4 pH 7.2)) and 30% sucrose solution for 1 hr)]. After overnight postfixation, brains were cryoprotected in increasing concentrations of sucrose solution (10%, 20%, and 30%). Brains were sectioned coronally on a freezing, sliding microtome at 40 μm, placed in 24 consecutive compartments containing cold cryoprotectant solution (30% sucrose and 30% ethylene glycol in PB), then stored at -20°C (Rosene et al., 1986).

### Histology

#### Immunohistochemistry (ICC)

In both Cohorts, every eighth section through the entire brain was processed using ICC for the relevant tracer. In Cohort 1, adjacent sections through the striatum were processed for calbindin-D28k protein (CaBP) in order to localize the position of anterogradely labeled fibers in striatal subregions (see *Amygdalo-striatal path* section in *<u>Cortico-amygdalo-striatal analyses</u>*; **Cohort 1**). In Cohort 2, CaBP-immunoreactivity (IR) in adjacent sections was used to localize tracer injections within striatal subregions (see *Injection site placement;* **Cohort 2**). For all experiments, tissue was rinsed in phosphate buffer with 0.3% Triton-X (PB-TX) overnight. The next day, brain slices were treated with an endogenous peroxidase inhibitor for 5 minutes, and then underwent more rinses in PB-TX. Sections were then pre-incubated for 30 minutes in 10% normal goat serum blocking solution with PB-TX (NGS-PB-TX). All sections were then incubated in primary antisera to LY (1:2000; Invitrogen, rabbit), FS (1:2000; Invitrogen, rabbit), FR (1:1000; Invitrogen, rabbit), WGA (1:2500-1:7500 depending on dilution curve; Sigma, rabbit), or CaBP (1:10000; Millipore Bioscience Research Reagents, mouse) at 4°C for four nights. Sections were then thoroughly rinsed, again blocked with 10% NGS-PB-TX, and incubated for 40 minutes in the appropriate biotinylated secondary antibody. After more rinses, sections with bound anti-LY, anti-FR, anti-FS, anti-WGA, or anti-CaBP antibodies were incubated in an avidin-biotin complex (Vectastain ABC kit; Vector Laboratories), and visualized with 3,3’-Diaminobenzidine (DAB), activated with 0.3% hydrogen peroxide (H_2_O_2_).

#### Cresyl Violet

In order to localize cortical cytoarchitectonic boundaries for each animal (both Cohorts 1 and 2), we stained 1:24 adjacent or near-adjacent sections for each case with cresyl violet (Chroma-Gesellschaft, West Germany).

#### Acetylcholinesterase (AChE)

In Cohort 1, AChE staining in adjacent sections was used to identify injection site location in the basal nucleus, while in Cohort 2, AChE staining was used to localize tracer-labeled neurons in specific basal nucleus subdivisions. For both, we stained 1:24 adjacent or near-adjacent sections for AChE, using the Geneser-Jensen technique (Geneser-Jensen and Blackstad, 1971).

#### Analysis

##### Cytoarchitectural criteria for the PFC and insula (Fig. 2)

The PFC is defined as the frontal lobe anterior to the arcuate sulcus. Within the PFC, we define the ventromedial and orbital subdivisions according to the nomenclature of Carmichael and Price (Carmichael and Price, 1994), and the dorsomedial (dmPFC), dorsolateral (dlPFC), and ventrolateral (vlPFC) subdivisions according to Petrides et al. (Petrides and Pandya, 1999, 2002). Agranular insula subdivisions were defined according to Carmichael and Price (Carmichael and Price, 1994), and dysgranular and granular insula subdivisions were defined according to Mesulam & Mufson (Mesulam and Mufson, 1982). Regardless of nomenclature, all cytoarchitectural maps of the PFC recognize the gradual transition from agranular to granular laminar organization in the basoventral and mediodorsal direction, based on laminar complexity and the relative development of layer IV (Brodmann, 1909; Preuss and Goldman-Rakic, 1991; Barbas, 2007; Nieuwenhuys et al., 2007). The insula similarly progresses from agranular to granular, but in a rostroventral to caudodorsal direction (Mesulam and Mufson, 1982). See Table 2 for descriptions of laminar characteristics of our cytoarchitectural divisions.

**Figure 2:**
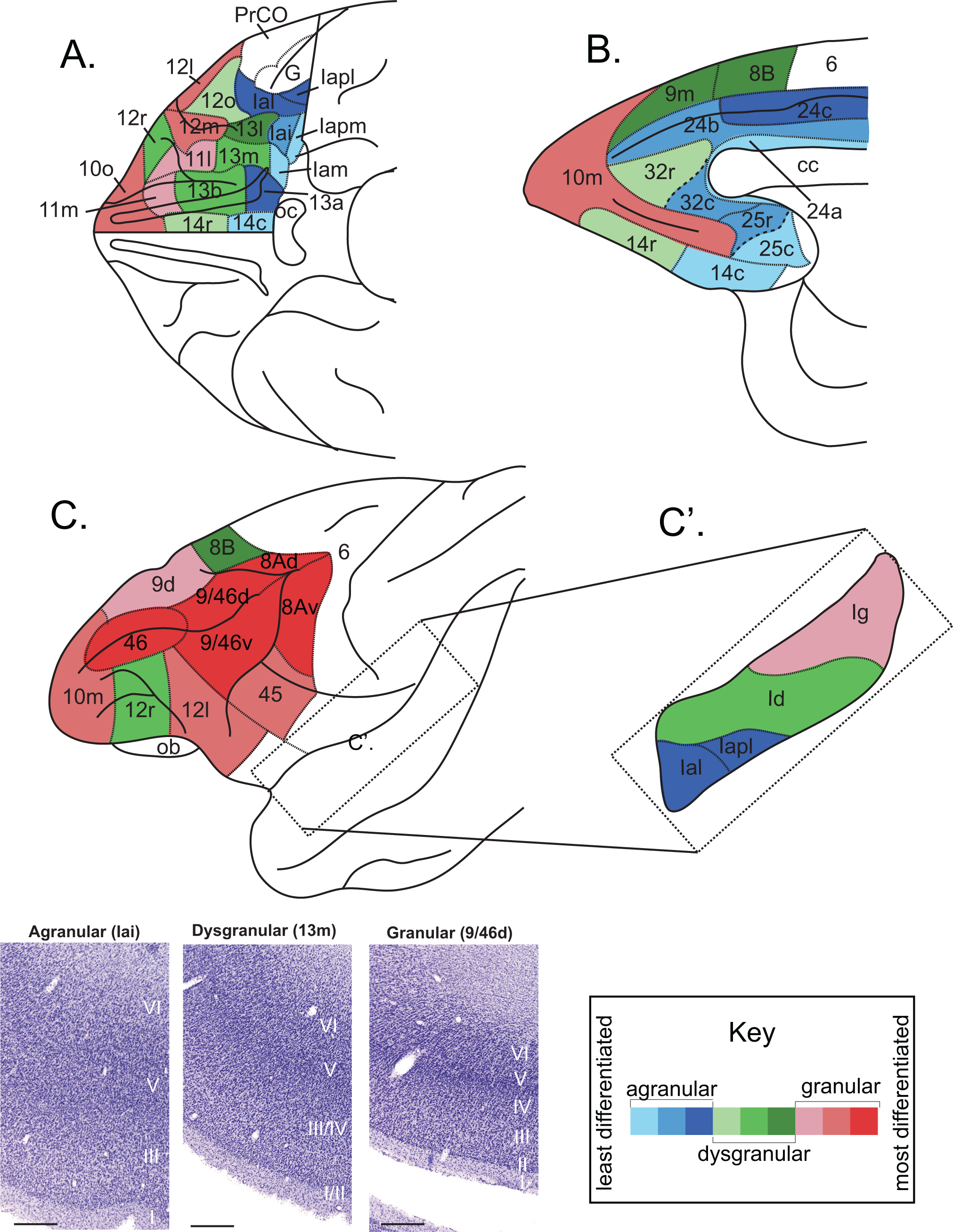
Relative levels of cortical lamination in the PFC and Insula in the Macaque. ***A***. Ventral view ***B***. Sagittal view from the midline ***C***. Lateral view, with lateral fissure ’opened’ for ***C’.*** view of insula. Image adapted from Carmichael & Price, 1994 and Saleem & Price, 2008. The key illustrates the range of cortical differentiation; shades of blue indicate agranular cortex, shades of green indicate dysgranular cortex, and shades of red indicate granular cortex. The darker the shade within each granularity grouping indicates increased development of layer IV, layer II, and/or layer V of cortex. Insets are from cresyl violet stained sections that give examples of laminar regions in agranular, dysgranular and granular cortices. Bar = 500 um. *Abbreviations: 6, area 6 (premotor/supplementary motor area); 8Ad, dorsal area 8A; 8Av, ventral area 8A; 8B, area 8B; 9d, dorsal area 9; 9m, medial area 9; 9/46d, dorsal area 9/46; 9/46v, ventral area 9/46; 10m, medial area 10; 10o, orbital area 10; 11l, lateral area 11; 11m, medial area 11; 12l, lateral area 12; 12m, medial area 12; 12o, orbital area 12; 12r, rostral area 12; 13a, area 13a; 13b, area 13b; 13l, lateral area 13; 13m, medial area 13; 14c, caudal area 14; 14r, rostral area 14; 24a, area 24a; 24b, area 24b; 24c, area 24c; 25c, caudal area 25; 25r, rostral area 25; 32c, caudal area 32; 32r, rostral area 32, 45, area 45; 46, area 46; 9/46d, dorsal area 9/46; area 9/46v, ventral area 9/46; cc, corpus callosum; G, gustatory cortex; Iai, intermediate agranular insula area; Ial, lateral agranular insula area; Iam, medial agranular insula area; Iapl, posterolateral agranular insula area; Iapm, posteromedial agranular insula area; Id, dysgranular insula; Ig, granular insula; ob, olfactory bulb; oc, optic chiasm; PrCO, precentral opercular area*.

**Table 2:** Laminar characteristics of the PFC and insula subdivisions, and 3- and 9-group granularity assignments. Each column indicates the main granularity grouping of every cortical area examined (i.e. the relative thickness of layer IV of cortex). The darker the shade within each sub-section within each column (i.e. ’Agranular 1’ vs. ’Agranular 2’) represents the increasing degree of layer V sublamination and/or layer IV development within each granularity assignment.

| Agranular |  |
| --- | --- |
| Granularity Grouping | Reason for granularity grouping |
| Agranular 1 |  |
| lam | No layer IV, and no layer V sublamination. (Carmichael & Price, 1994) |
| lapm | No layer IV, and no layer V sublamination. (Carmichael & Price, 1994) |
| 25c | No layer IV, less developed compared to 32c, 32r, and 25r. (Carmichael & Price, 1994) |
| 24a | Has only 4 cortical layers and rudimentary lamination. (Carmichael & Price, 1994) |
| 14c | Has only 4 cortical layers. (Carmichael & Price, 1994) |
| Agranular 2 |  |
| 32c | No layer IV; Caudal 32 resembles 25r; has horizontal striations of pyramidal cells in layer V. (Carmichael & Price, 1994) |
| 25r | No layer IV; caudal 32 resembles 25r. (Carmichael & Price, 1994) |
| 24b | No layer IV; has more prominent vertical striations in layer V compared to 24c. (Carmichael & Price, 1994) |
| lai | No layer IV; has partially sublaminated layer V; superficial layer V has row of large pyramidal cells. (Carmichael & Price, 1994) |
| Agranular 3 |  |
| lal | No layer IV; layer V is sublaminated. (Carmichael & Price, 1994) |
| lapl | No layer IV; layer V is sublaminated. (Carmichael & Price, 1994) |
| 13a | No layer IV; layer V is sublaminated. (Carmichael & Price, 1994) |
| 24c | No layer IV; Has pyramidal cell aggregates in layer V. (Carmichael & Price, 1994) |

| Dysgranular |  |
| --- | --- |
| Granularity Grouping | Reason for granularity grouping |
| Dysgranular 1 |  |
| 32r | Is more developed than 32c and 25r and is slightly dysgranular (Carmichael & Price, 1994). No layer V sublamination. |
| 12o | Very weakly staining layer IV, no layer V sublamination. (Carmichael & Price, 1994) |
| 14r | Has distinct small cells in layer II, no horizontal or vertical striations in layer V. (Carmichael & Price, 1994) |
| Dysgranular 2 |  |
| 13b | Has horizontal and vertical striations in layer V. (Carmichael & Price, 1994) |
| 12r | Has vertical striations in layer V, but it is not sublaminated. (Carmichael & Price, 1994) |
| 13m | Has a sparse granular layer. Has no/less distinct sublamination of layer V (and layer III). (Carmichael & Price, 1994) |
| ld | Has thin layer IV with clusters of granular cells interrupted by pyramidal cells. No layer II. Almost identical in appearance to dysgranular component of caudal OFC. (Mesulam & Mufson, 1982) |
| Dysgranular 3 |  |
| 13l | Has poorly developed layer IV and has clearly sublaminated layer V. (Carmichael & Price, 1994) |
| 8B | Layer II is well defined; Layer IV is poorly developed and layer V blends with layer VI, but layer V is sublaminated (it has a few darkly stained cells in layer Va). (Petrides & Pandya, 1999) |
| 9m | Has a more narrow and cell sparse layer IV than 9. (Petrides & Pandya, 1999) |

| Granular |  |
| --- | --- |
| Granularity Grouping | Reason for granularity grouping |
| Granular 1 |  |
| 9d | Has a narrow layer IV. Layer Va has deeply pigmented cells and layer Vb blends with layer VI. (Petrides & Pandya, 1999) |
| lg | Layer II has moderately advanced granularity, and layer IV has prominent granularity. Layer V and VI are not fully differentiated from each other; (Layer III is bilaminar). (Mesulam & Mufson, 1982) |
| 11m | Granular with bilaminar layer V. Layer V is thinner and more trilaminar than in 13m. Outer and inner sublaminae are broken into aggregates of neurons. (Carmichael & Price, 1994) |
| 11l | Granular with bilaminar layer V containing aggregates of pyramidal cells. Layer V is thinner and more trilaminar than in 11m. Does not have outer and inner sublaminae broken up into neuron aggregates. (Carmichael & Price, 1994) |
| Granular 2 |  |
| 10o | Is more granular than 11m and 11l, but layer IV not as clear. Layer V is thinner than in 11m or 11l; contains vertical and horizontal striations in layer V, and layer V is thinner than 11l and 11m. (Carmichael & Price, 1994) |
| 10m | Has completely developed II and IV; has horizontal and vertical striations in layer V. Layer IV is more granular than 11m and 11l but not well demarcated, thin layer V with radial striations. (Carmichael & Price, 1994) |
| 12m | Has completely developed II and IV; has clearly sublaminated layer V. Is more granular than 12r, 13l, and 12o. (Carmichael & Price, 1994) |
| 12l | Has completely developed layers II and IV; has prominent/sharp sublamination of layer V. (Carmichael & Price, 1994) |
| 45 | Well developed layer IV with medium sized layer V (also has distinct and uniquely thick layer III). (Petrides & Pandya, 2001) |
| Granular 3 |  |
| 8Av | Layer IV is broad and densely packed with small neurons (Layer II is also developed and densely packed). Layer V is less dense, but is bilaminar. Resembles 8Ad, but layer IV is broader, and layer III pyramidal neurons are more densely packed. (Petrides & Pandya, 1999) |
| 8Ad | Layer IV is broad and densely packed with small neurons (Layer II is also developed and densely packed). Layer V is less dense, but is bilaminar. Resembles 8Av, but layer IV is less broad, and layer III pyramidal neurons are less densely packed. (Petrides & Pandya, 1999) |
| 46 | Layer IV is well developed; layer V is sublaminated (layer Va has small number of deeply stained cells and layer Vb blends with layer VI). (Petrides & Pandya, 1999) |
| 9/46d | Similar features as 46, but layer III has large and darkly stained pyramidal neurons; Located dorsally. (Petrides & Pandya, 1999) |
| 9/46v | Similar features as 46, but layer III has large and darkly stained pyramidal neurons; Located ventrally. (Petrides & Pandya, 1999) |

##### Categorization of cortical regions by architecture

For quantitative analysis, we categorized cortical areas into either 3 (general) or 9 (refined) categories of cortical hierarchy, based on examining layer IV thickness (for 3 category granularity) and the extent of layer V sublamination (for 9 category granularity) under Nissl stain (Carmichael and Price, 1994) (Table 2). For 3 category granularity, cortical regions were categorized as agranular cortex if they did not have layer IV, dysgranular if they had incompletely developed layer IV, and granular if they had completely developed layer IV.

#### Cohort 1

##### Injection site placement

Relative injection site placement within the Bmc, Bi, and Bpc was determined using adjacent AChE sections for each animal. Throughout the rostrocaudal extent of the basal nucleus, the Bmc is darkly stained with AChE, the Bi has intermediate AChE staining, and the Bpc is lightly stained with AChE, corresponding to the cytoarchitectural gradients in the nucleus (Amaral and Bassett, 1989). All nuclei, and specifically basal nucleus subdivisions, were traced along with landmarks, such as blood vessels across sections, and aligned over tracer-labeled sections in Adobe Illustrator Creative Suite (CS).

Cases with tracer leakage into overlying structures or white matter tracts were not included in the analysis. To confirm our assessment of injection site placement, we also examined the pattern of retrograde labeling in brain regions known to be a source of afferents to each subdivision. For example, the Bmc receives inputs from the inferotemporal cortex (Herzog and Van Hoesen, 1976), while the Bpc does not; the Bpc receives inputs from the hippocampus in contrast to the Bi and Bmc (Saunders et al., 1988; Fudge et al., 2012).

#### Cortico-amygdalo-striatal analyses

##### Cortico-amygdala path

Retrogradely labeled cells in 1:24 sections were charted through the entire rostral-caudal extent of the PFC and insula using an Olympus AX70 microscope interfaced with Neurolucida via a video CCD (Microbrightfield, Williston, VT). We determined cytoarchitectural boundaries of cortex on adjacent Cresyl violet-stained sections under the microscope, based on criteria described above. These labeled charts were then aligned onto maps of retrogradely labeled cells in Adobe Illustrator CS. The number of retrogradely labeled cells in each cortical subdivision was counted using the Adobe Illustrator “objects” count feature. Cortical subregions were then categorized into 3- and 9-category “agranular”, “dysgranular”, and “granular” groupings as described above (Table 2).

After assignment of each cortical area to a cortical granularity category, we used the “*circlize”* package in R (R Foundation for Statistical Computing, Vienna, Austria) to generate chord diagrams of our retrograde cortical data, using our 3 and 9 category classifications of cortical granularity (Gu et al., 2014). Chord diagrams are an effective way to visualize the directionality and connectivity between different nodes of a large data set. ‘Fragments’, which represent each node, are found along the entire perimeter of the chord diagram. The small tic marks and numbers for each ‘fragment’ indicate number of labeled cells. Chord diagrams are organized such that regions containing retrogradely labeled cells (i.e. ‘agranular’ cortex, ‘dysgranular’ cortex, etc.) are located at the top of the diagram as individual ‘fragments’, and injection site locations (i.e. ‘J12LY’, ‘J15LY’, etc.) are located at the bottom of the diagram as individual ‘fragments’. This organization thus reflects the number of retrogradely labeled cells (top axis) associated with specific retrograde injection sites (bottom axis), and the quantitative relationships between the ’strength’ of the pathway (number of labeled cells) across cases.

##### Amygdalo-striatal path

Anterogradely labeled terminal fibers were hand-charted through the rostro-caudal extent of the striatum with the aid of a drawing tube under dark-field illumination. Labeled thick fibers that did not contain boutons were considered fibers of passage and were therefore not included. All hand-drawn charts were then scanned at high resolution and converted in Adobe Illustrator CS for formatting. Striatal boundaries were then determined through comparison to adjacent or near-adjacent CaBP-labeled sections. CaBP-labeled sections were projected onto paper maps of anterogradely labeled fibers in the striatum. Landmarks including blood vessels and fiber tracts were carefully aligned, and CaBP-positive and CaBP-negative areas of the striatum were drawn in. Maps were then scanned into digital format at high resolution, and converted in Adobe Illustrator CS.

#### Cohort 2

##### Injection site placement

Injection sites were targeted to the rostral ventral striatum, and also to regions of the ventromedial striatum posterior to the anterior commissure, based on anterograde maps from Cohort 1. Injection site position was determined with reference to adjacent CaBP-labeled sections. In rostral ventral striatal regions, CaBP-poor regions were used to identify the ‘shell’, while CaBP-rich regions in the rostral ventral striatum were used to identify the ’core’ (Meredith et al., 1996; Fudge and Haber, 2002). Collectively, we classify the ‘shell’ and ‘core’ as ‘classic’ ventral striatum. All ventral striatal regions caudal to the level of the ‘shell’ and ‘core’ are referred to as ‘extended’ caudal ventral striatum (Fudge and Haber, 2002). All injection sites that resulted in retrogradely labeled cells in the amygdala were used for cortico-‘limbic’ striatum and amygdalo-striatal analyses.

#### Amygdalo-striatal path

The location and quantification of retrogradely labeled cells in basal nucleus subdivisions was mapped in 1:24 sections, with reference to adjacent AChE stained sections. All retrogradely labeled cells in the amygdala were subsequently sorted by subdivision (Bmc, Bi, and Bpc) based on levels of AChE activity and cellular features, and analyzed with respect to position of the injection within the striatum.

#### Cortico-‘limbic’ striatal path

Only injection sites resulting in significant numbers of labeled cells in the basal nucleus (>10 cells) were used for these analyses. We quantified retrogradely labeled cells in the entire PFC and insula, and applied the same criteria for classification, analysis, and visualization described for Cohort 1.

#### Comparing cortico-amygdala-striatal and cortico-striatal pathways

As a way of estimating whether amygdala neuronal populations projecting to different striatal regions received a similar balance of agranular, dysgranular, and granular cortical inputs, we used a ‘ratio of ratios’ approach to examine data across Cohort 1 (cortico-amygdala) and Cohort 2 (amygdala-striatal). While this analysis does not directly examine cortical inputs to amygdala-striatal projections in the same animal, our goal was to examine the proportions of cortical inputs associated with each basal nucleus subdivision. We first pooled the sum of all agranular, dysgranular, and granular labeled cell counts resulting from Bmc, Bi, and Bpc injection sites (i.e. cortico-amygdala data), and converted these sums into percentages. Only basal nucleus injection sites that were wholly confined to one of these subdivisions were used. Once the proportion of labeled neurons in the agranular, dysgranular, and granular cortex were determined for Bmc, Bi, and Bpc injection groupings, we multiplied these percentage values by the number of retrogradely labeled cells in Bmc, Bi, and Bpc resulting from each striatal injection site in Cohort 2. We then converted this sum into a final percentage value to assess the balance of agranular, dysgranular, and granular cortical inputs to amygdalo-striatal projecting cells for each striatal site. We conducted bootstrap analysis to check the ‘stability’ of our final percentage values in ‘indirect’ pathway calculations. We did 4 replicates of the data, and found that the average difference of full versus bootstrap data was = 1.4%, the median was 1.0%, the standard deviation was 0.01198, the maximum was 5.6%, and the minimum was 0.1%. This was deemed acceptable. The proportion of labeled cells in the agranular, dysgranular, and granular cortices for the indirect pathway (conjunction of Cohort 1 cortico-amygdala data and Cohort 2 amygdalo-striatal data) were then compared to the proportion of labeled cells in agranular, dysgranular, and granular cortices for each injection site associated with the direct cortico-striatal pathway (Cohort 2). Results were expressed as the proportion of labeled neurons in the agranular, dysgranular, and granular cortex associated with each path for each striatal injection site. Sample variance for each granularity grouping and both pathway types was collected and compared.

#### Cluster analysis for cortico-amygdala and cortico-striatal data

In order to identify cortical subregion connectivity ‘fingerprints’ of specific basal nucleus and striatal subregions in a quantitative way, we also used k-means cluster analysis on whole cortico-amygdala and whole cortico-striatal retrograde data. K-means cluster analysis is an unsupervised machine learning technique that classifies variables (i.e. retrogradely labeled cells in cortical subregions) into ‘*k*’ groups such that variables with similar characteristics cluster together, while dissimilar variables are clustered in non-overlapping, separate groups (MacQueen, 1967; Kaufman and Rousseeuw, 2009).

We organized the data matrix rows as we did our chord diagram data, i.e. by ascending granularity grouping (see Table 2) and with cortico-amygdala data matrix columns organized from caudo-ventral to rostro-dorsal injection site location, and cortico-striatal data matrix columns from rostral to caudal injection site location. Cell counts in cortical areas for each injection site were z-scored (using the formula (x - μ) / σ for each injection site, where x is the number of cells in a specific cortical subregion, μ is the mean cell count of all cortical subregions examined for a specific injection site, and σ is the cell count sample standard deviation of all cortical subregions examined for a specific injection site). Z-scoring (or scaling in general) normalizes the data, which accounts for differences in variance across data points, ensures that cluster plots are not biased by either very high or very low cell count totals, and allows all variables to be considered with equal importance.

In order to approach our clustering in an unbiased manner, and to ensure we were not under- or over-clustering data, we used the ‘silhouette method’ as a guide to determine the optimal number of ‘*k*’-clusters to use per data set (Rousseeuw, 1987). The average silhouette width measures how well each variable fits within a cluster; whichever value of *k* is closest to a value of 1 average silhouette width represents high intra-group similarity and high inter-group difference, and was selected as the ‘*k*’ value for that data set. We selected the largest value of *k* that was greater than a 0.5 average silhouette width score.

For all of these analyses, we used the ‘tidyverse’, ‘cluster’, and ‘factoextra’ packages in R Studio. We used the ‘set.seed’ function to ensure stability and repeatability of our data simulations. We also set ‘n.start’ to 25, meaning that 25 iterations of our code are run, and the best cluster formatting that results in the lowest within-cluster variation is selected. Datasets and coding information for all data on are deposited on https://figshare.com/s/2ad6809dccb896995407.

In order to interpret each cluster, we inspected the patterns of z-scores for each cortical area associated with each cluster. Z-scores were ranked ‘high’ (mean z-score between 1.5001 – 5.5000), ‘mean/above mean’ (mean z-score between 0.0000 – 1.5000), ‘slightly below mean’ (mean z-score between -0.0999 - -0.0001), or ‘largely below mean’ (mean z-score between -0.6000 - -0.1000), based on the distribution of z scores for all cortical regions around the mean. This was done for each pathway. Descriptive names were created for each cluster according to its unique identity.

## Results

### Cortico-amygdalo-striatal paths (Cohort 1)

#### Basal nucleus injection site placement (Fig. 3)

In the Bmc, there were three injections placed at slightly different levels—cases J12LY, J16LY, J12FR—as well as five injections at different levels of the Bpc—cases J20LY, J15LY, and J14FR in rostral Bpc, and cases J15FS and J14FS in caudal Bpc, all reported previously (Cho et al., 2013). Three new injections at slightly different levels of the Bi—cases J47FS, J44FR, and J52LY were made and assessed for a more complete survey of the basal nucleus subdivisions. The Bi injection in J47FS was the most ventral, the injection in J52LY extended slightly more dorsally, and the injection in J44FR was most dorsal and lateral, straddling the border with the lateral Bmc.

**Figure 3:**
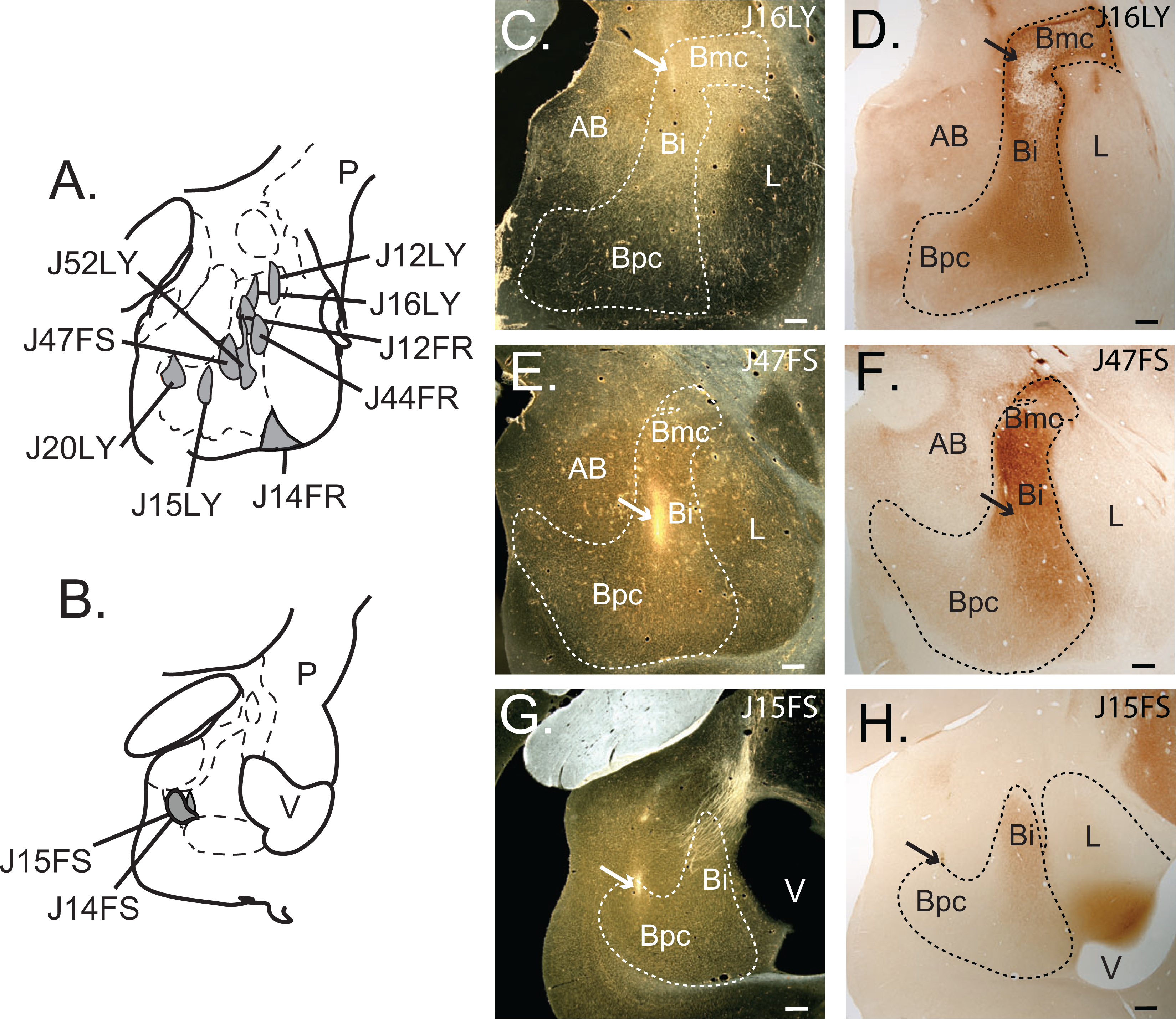
Schematic of injection site locations in basal nucleus. ***A***. Injection site locations (gray) in rostro-central Bmc, Bi, and Bpc. ***B***. Injection site locations (gray) in caudal Bpc. ***C***. Dark-field photomicrograph of injection site location for J16LY, indicated with a white arrow. ***D***. Adjacent AChE-stained section matched to ***C.*** to localize injection to basal nucleus subdivision (Bmc). ***E***. Dark-field photomicrograph of the injection site location for J47FS, indicated with a white arrow. ***F***. Adjacent AChE-stained section matched to ***E.*** to localize injection site within basal nucleus subdivision (Bi). ***G.*** Dark-field photomicrograph of the injection site location for J15FS, indicated with a white arrow. ***H.*** Adjacent AChE-stained section matched to ***H.*** to localize injection site within basal nucleus subdivision (Bpc). Photos taken at 2x magnification. Scale bars = 500 μm. *Abbreviations: AB, accessory basal nucleus; Bi, intermediate basal nucleus; Bmc, magnocellular basal nucleus; Bpc, parvicellular basal nucleus; L, lateral nucleus; P, putamen; V, ventricle*.

#### Cortical inputs along basal nucleus subdivisions (Fig. 4)

After all injections in the basal nucleus, a general pattern emerged in which injection sites placed in the Bpc resulted in many labeled cells in the PFC and insula, restricted to the agranular PFC and insula, while increasingly dorsal and rostral injections in the Bpc, Bi, and Bmc resulted in labeled cells in broader cortical regions, as detailed below.

**Figure 4:**
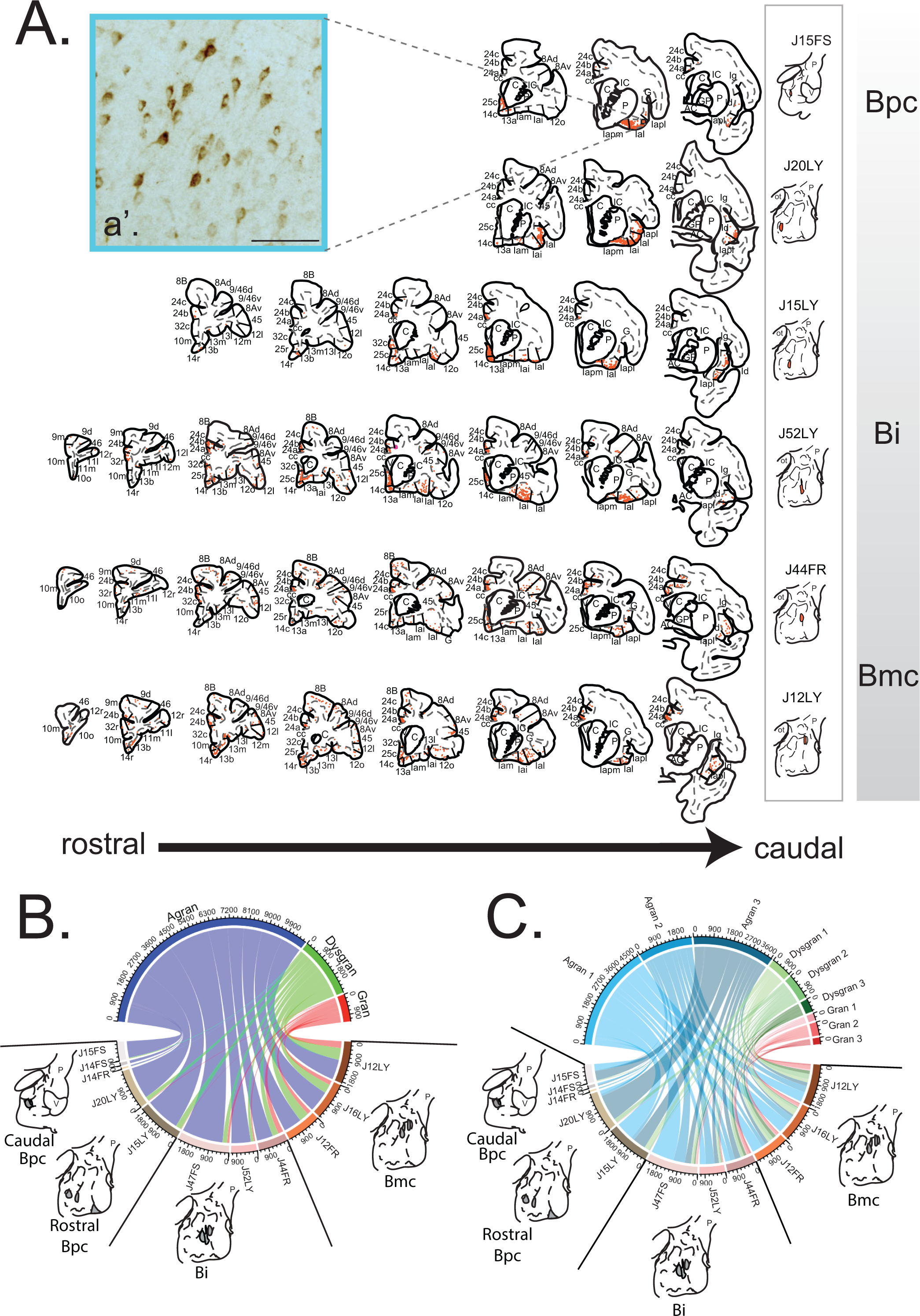
Cortico-amygdala path. ***A.*** Representative charts showing the pattern of retrogradely labeled cells in the PFC and insula resulting from 6 non-overlapping injection sites along the dorsal-ventral extent of the basal nucleus. ***a’*.** The photomicrograph inset shows an example of densely concentrated retrogradely labeled cells in the Ial of J15FS. Scale bar = 100 μm ***B.*** Chord diagram showing 3-group quantitative analysis of all retrogradely labeled cells (1:24 sections) through PFC and insula, classified by laminar differentiation. The top axis of this diagram shows the total number of labeled cells in agranular (blue), dysgranular (green), and granular (red) cortical areas across all cases examined. The bottom axis shows the number of labeled cells in agranular, dysgranular, and granular cortices resulting from each basal nucleus injection site. Injection sites are arranged counter-clockwise from the most caudal-ventral (left) to most rostral-dorsal injection site location (right). 1 tick mark= 180 cells. ***C.*** Chord diagram showing 9-group granularity analysis. The top axis shows the total number of labeled cells in agranular (3 shades of blue), dysgranular (3 shades of green), and granular (3 shades of red) cortical areas across all cases examined. The darker the shade within each granularity grouping indicates increased development of layer IV and/or layer V per Table 2. 1 tick mark represents 180 cells. *Abbreviations: AC, anterior commissure; C, caudate; cc, corpus callosum; GP, globus pallidus; IC, internal capsule; oc, optic chiasm; P, putamen, V, ventricle. For cortical abbreviations, see “Abbreviations” in* Figure 2 *legend*.

#### Bpc

Injection sites in the Bpc (cases J15FS, J14FS, J14FR, J20LY, J15LY) resulted in labeled cells largely confined to agranular PFC, i.e. area 25c, 14c, and agranular insula areas (Fig. 4A). J15FS and J15LY had the most labeled cells in agranular insula subdivisions Iapm and Ial, J14FS had most labeling in Iai and Iapl, J14FR had most labeling in Iapl, and J20LY had most labeling in Ial and Iapl. Across Bpc cases, the majority of labeled cells was in the agranular insula rather than PFC, however, the proportion of labeled cells in the agranular PFC increased gradually as the injection sites were positioned more laterally. Case J15LY, which is an injection in the ‘transition’ between the Bpc and Bi, had additional labeling in agranular area 24a. J20LY had relatively more labeled cells in dysgranular insula compared to other Bpc sites.

#### Bi

Bi injections sites (cases J47FS, J52LY, and J44FR) resulted in the majority of labeled cells in agranular cortices, but with a relatively greater contribution of labeled cells in dysgranular and granular cortices compared to Bpc (Fig. 4A). Similar to Bpc, high numbers of labeled cells were found in 25c, 14c, and agranular insula. Case J47FS had the most labeled cells in Iapm and Iai, case J52LY had the majority of labeled cells in Iai and Ial, and case J44FR had the most labeled cells in Iapl. In contrast to Bpc sites, a broader distribution of labeled cells occupied relatively more differentiated agranular areas, including 32c, 24a, 24b, and 13a, as well as labeled cells in dysgranular regions such as area 13b. J44FR, located in a ‘transition’ region between Bi and Bmc, had relatively more labeled cells in agranular area 24c, dysgranular insula, and 8B, in addition to labeled cells specifically in granular areas 12l and 8Ad.

#### Bmc

While the majority of labeled cells remained in the agranular regions of PFC and insula, injections in Bmc cases (J12FR, J16LY, and J12LY) had the broadest distribution of labeled cells. Similar to Bpc and Bi, Bmc had many labeled cells in agranular areas 25c, 14c, and agranular insula. Similar to Bi cases, Bmc cases had high levels of labeled cells in agranular areas 24a, 24b, and 24c, but in contrast, also had moderate numbers of labeled cells in dysgranular areas 14r, 8B, 13m, and 12o, and some labeled cells in granular areas 45, 9, 8Ad, 8Av, and 46.

Quantitative analyses revealed that cortical granularity subtype, rather than association with the insula or PFC, predicted inputs across, and even within, basal nucleus subdivisions (Fig. 4B). The agranular cortices in both PFC and insula contained labeled cells after all basal nucleus injections, but were the sole contributor when injection sites were placed in the caudomedial Bpc. Labeled cells in dysgranular cortices appeared and increased in numbers following injections along the rostro-dorsal axis, from the rostro-medial Bpc to Bi to Bmc. Labeled cells in the granular cortices of the insula and PFC formed a relatively smaller contribution, seen only after injections in dorsal Bi and Bmc. The pattern of inputs was nested such that labeled cells in incrementally higher levels of cortical organization only appeared in conjunction with labeled cells in less differentiated cortical areas. Analyses using the more refined (9 category) granularity classification showed a similar, nested pattern (Fig. 4C). In these, subcategories of differentiation within a general granularity ‘level’ were themselves overlapping in a hierarchical manner.

#### Basal nucleus-striatal path

The distribution and relative density of anterogradely labeled fibers in the striatum from each basal nucleus injection site were mapped (Fig. 5). While all injections sites in the basal nucleus subdivisions resulted in labeled fibers in the shell of the nucleus accumbens, the distribution of labeled fibers varied predictably, expanding its distribution in the striatum as injection site position moved from the caudomedial to rostro-dorsal basal nucleus.

**Figure 5:**
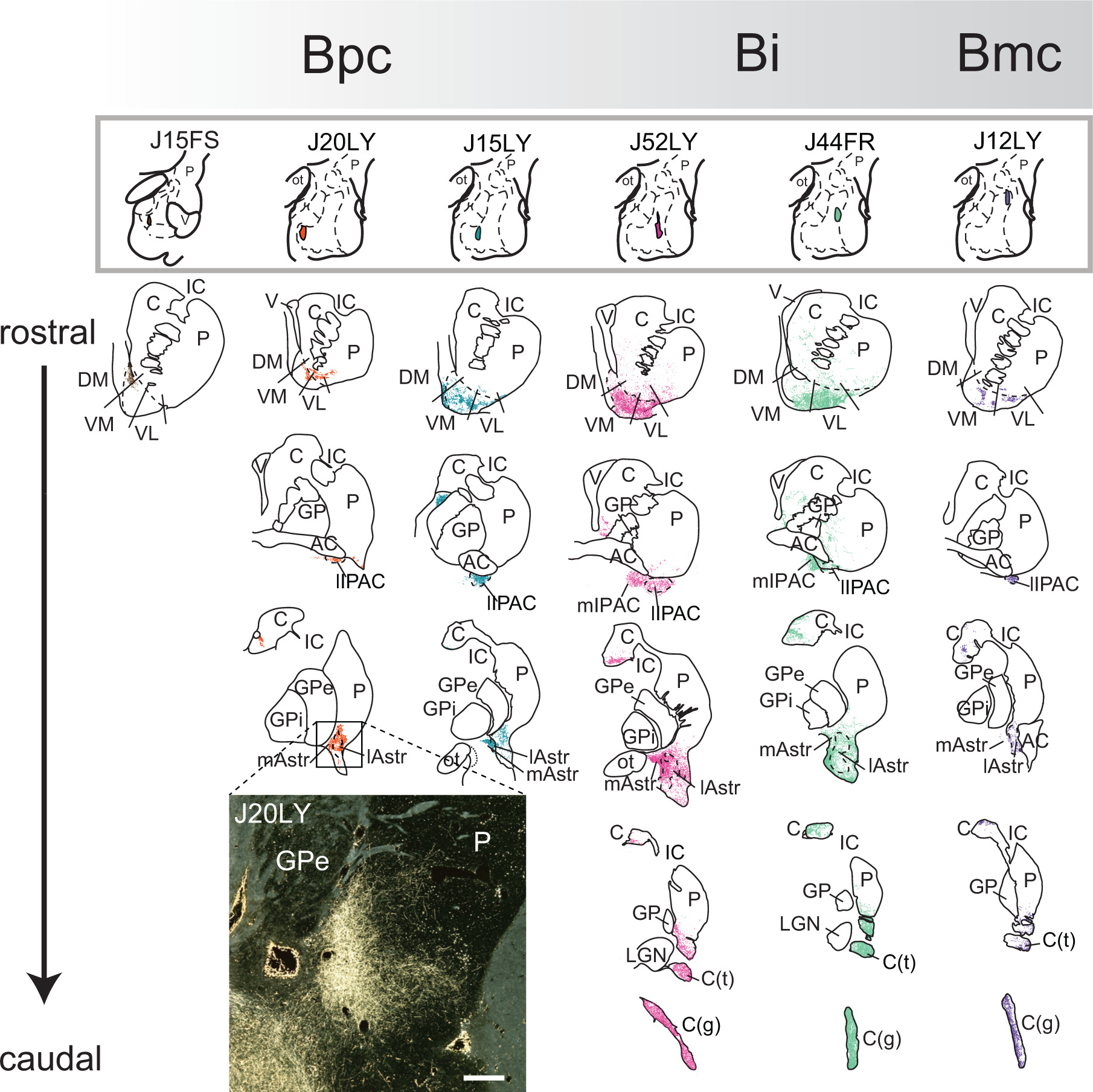
Representative charts showing distribution of anterogradely labeled fibers resulting from the same 6 non-overlapping injection sites shown in Fig. 4, along the dorsal-ventral extent of the basal nucleus. Injection sites arranged from caudo-medial (Bpc) to rostro-dorsal (Bmc) across the top. Striatal sections are organized from rostral to caudal levels under each injection site. Photomicrograph inset shows high density patch of labeled fibers in the caudal ventral putamen (darkfield). Bar = 500 μm. *Abbreviations: AC, anterior commissure; C, caudate; C(g), genu of the caudate nucleus; C(t), tail of the caudate nucleus; GP, globus pallidus; GPe; external globus pallidus; GPi, internal globus pallidus; IC, internal capsule; lAstr, lateral amygdalostriatal area; LGN, lateral geniculate nucleus; lIPAC, lateral interstitial nucleus of the posterior limb of the anterior commissure; mAstr, medial amygdalostriatal area; mIPAC, medial interstitial nucleus of the posterior limb of the anterior commissure; OT, optic tract; P, putamen; V, ventricle*.

#### Bpc

Caudomedial Bpc injections (Cases J15FS, J14FS) had labeled fibers largely confined to the dorsomedial shell of the ventral striatum, while more rostral and lateral Bpc injections (J14FR, J20LY, J15LY) resulted in additional labeled fibers in the medial and lateral shell of the ventral striatum, interstitial nucleus of the anterior commissure (IPAC, or fundus striatii), and a small region of the caudomedial putamen caudal to the anterior commissure.

#### Bi

Bi injection sites (cases J47FS, J52LY, and J44FR) resulted in labeled fibers in the same regions as Bpc inputs, except for the dorsomedial shell (Fig. 5). Compared to Bpc cases, Bi injection sites had additional light distributions of labeled fibers in the central rostral ’core’ of the ventral striatum, and moderate to heavy labeled fibers in the ventral body of the caudate nucleus, amygdalostriatal area, ventral putamen posterior to the anterior commissure, and tail of the caudate nucleus. Anterogradely labeled fibers continued caudally to fill the genu of the caudate nucleus.

#### Bmc

The broad pattern of labeled fibers (cases J12FR, J16LY, and J12LY) resembled those in the Bi, including high densities of labeled fibers in the ventral body of the caudate nucleus, amygdalostriatal area, ventral putamen posterior to the anterior commissure, and tail of the caudate nucleus. Labeled fibers in the rostral ventral striatum were mainly found in the lateral shell, and Bmc injections had more labeled fibers in central domains of the caudate head compared with Bi injections.

#### Cortico-striatal and amygdalo-striatal paths (Cohort 2)

##### Injection site placement

8 injections were placed into a range of rostral-caudal striatal regions that received amygdala input based on Cohort 1 anterograde data (Fig. 6). 6 sites had large numbers of retrogradely labeled cells in the basal nucleus, and were used for analysis. Of these, 2 injections are located in the rostral ventral striatum; J24WGA was placed in the CABP-negative dorsomedial shell, and J13WGA was placed in the CaBP-positive ventral striatal core. 4 injections are located in more caudal ventral striatal regions; J8WGA, J12WGA, and J11WGA were all placed at different levels of the caudoventral putamen. J41WGA was placed in the ventromedial body of the caudate nucleus posterior to the anterior commissure. 2 injections had relatively few labeled cells, and were classified as control cases (J35FR and J42WGA).

**Figure 6:**
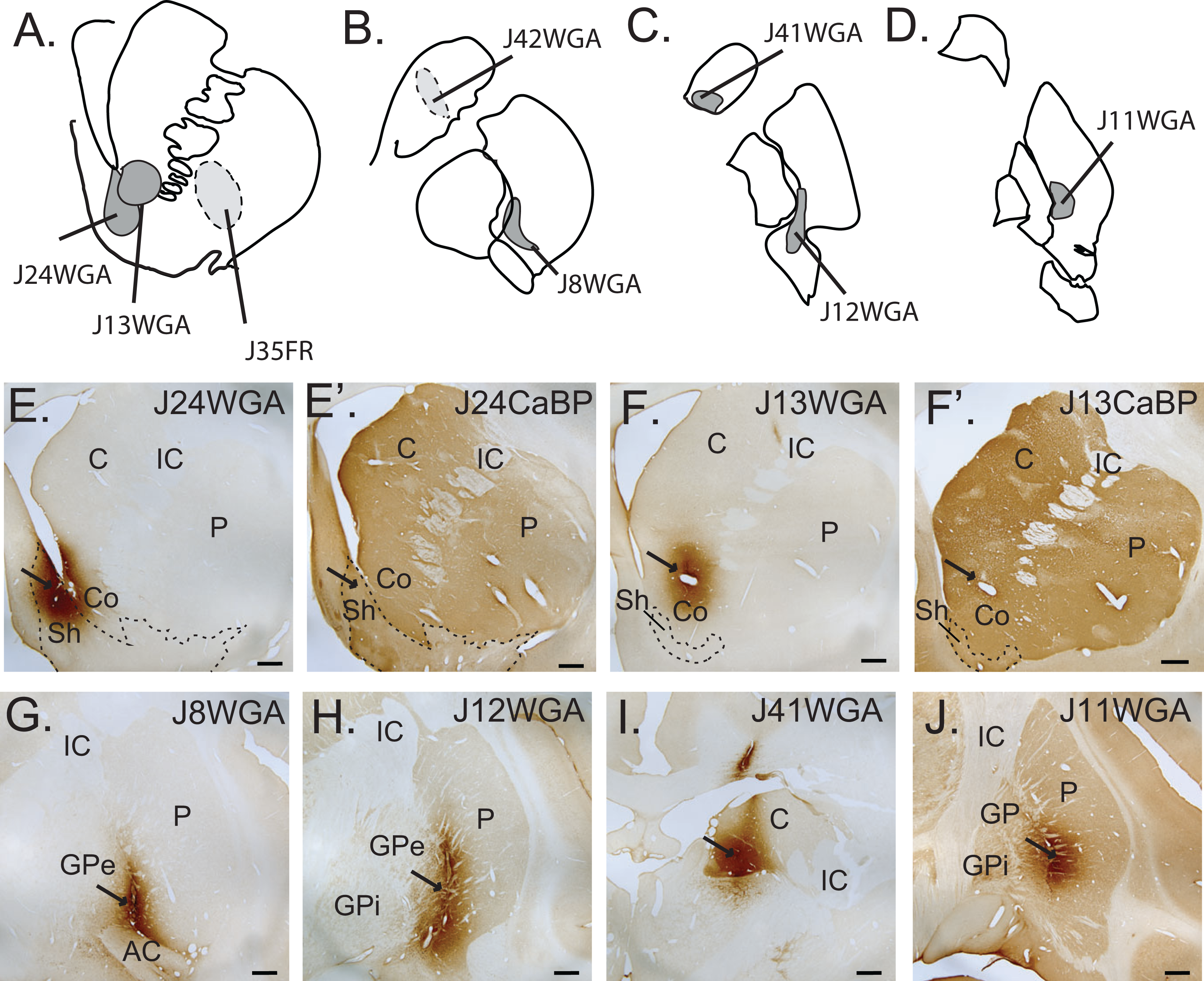
Striatal Injection sites in Cohort 2. ***A. - D.*** Schematics of retrograde injection site locations from rostral (A) to progressively caudal regions (B-D). Medium gray sites (solid lines) depict injections with significant labeled cells in amygdala, included for analysis; light gray sites (dotted lines) resulted in few labeled cells in the amygdala. ***E-F*.** Brightfield photomicrographs in the classic ventral striatum. ***E***. Injection site location in dorsomedial shell, with adjacent CaBP-stained section (***E’.***) showing injection alignment in CaBP-negative shell**. *F*.** Injection site in the rostral ’core’, with adjacent CaBP-stained section (**F’**.) showing shell/core boundary. ***G-J***. Caudal ’limbic’ striatum injection sites. ***G*.** caudoventral putamen at the level of the anterior commissure, ***H.*** caudoventral putamen posterior to the anterior commissure. ***I.*** ventral body of caudate nucleus**. *J.*** Injection site location in caudomedial putamen at the level of the hippocampus. Scale bars = 1mm. *Abbreviations: AC, anterior commissure; C, caudate; Co, ventral striatum core; GPe; external globus pallidus; GPi, internal globus pallidus; IC, internal capsule; P, putamen; Sh, ventral striatum shell*.

##### Amygdala inputs delineate ‘limbic’ striatum regions

The basal nucleus had many labeled cells after all injections (Figs. 7A). (Other amygdala nuclei, such as the accessory basal nucleus, had labeled cells in some cases (data not shown), in line with findings from previously published work (Fudge et al., 2002; Cho et al., 2013; Sharma et al., 2020).) In general, the total number of labeled cells in the basal nucleus was highest following rostral ventral striatal injections, decreasing with injections in progressively caudal sites (Fig 7B). With increasingly caudal injection sites, labeled cells also were found in the Bi and Bmc. The pattern of inputs from the basal nucleus subdivisions resembled the hierarchical layering of cortical inputs to the basal nucleus found in Cohort 1. The Bpc formed ubiquitous input to all striatal regions, with additional inputs sequentially added from Bi and Bmc, respectively, in increasingly caudal ventral striatal regions. We termed these caudal regions ‘extended’ caudal ventral striatum.

**Figure 7:**
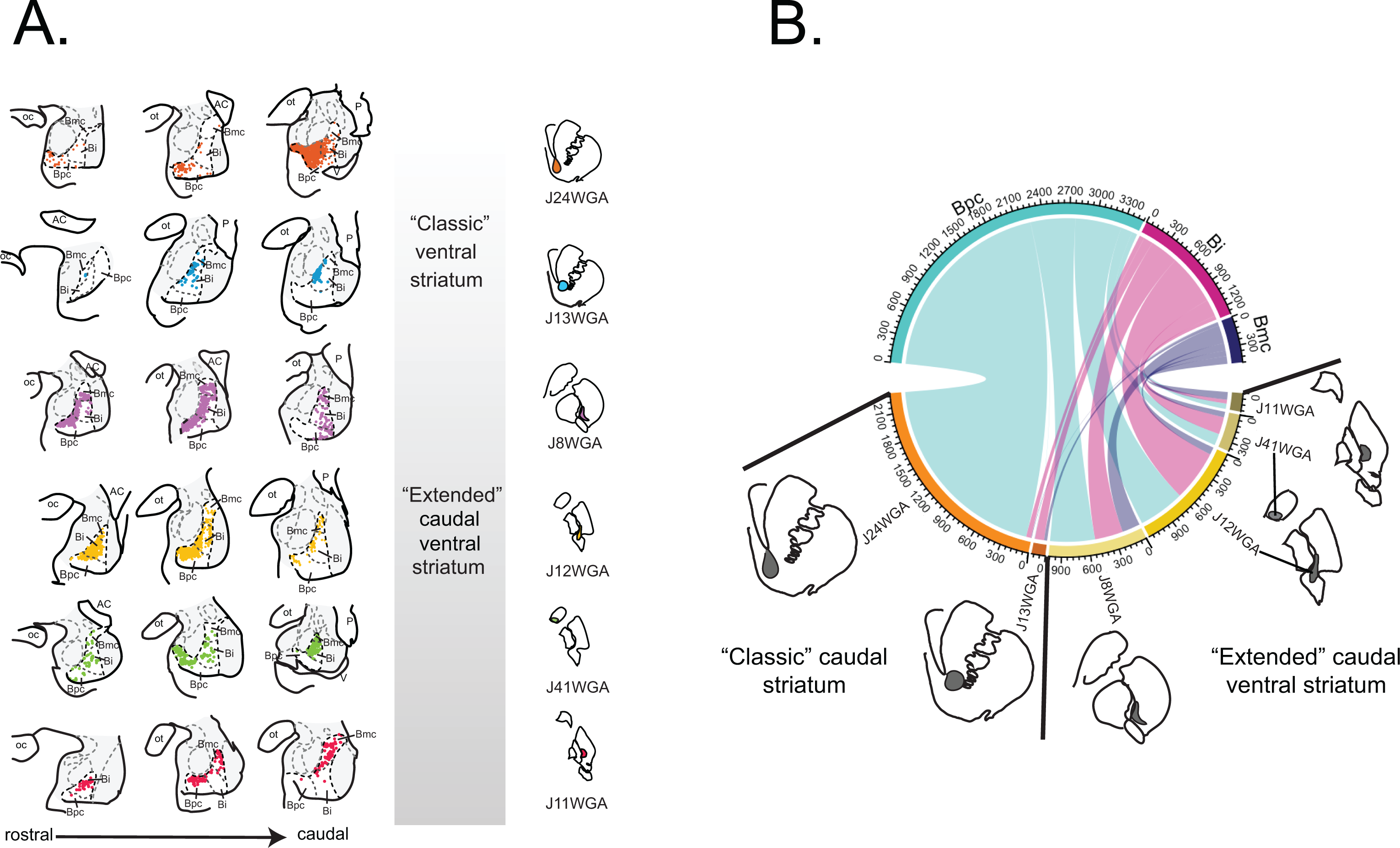
Amygdalo-striatal path. ***A.*** Charts of retrogradely labeled cells through the rostrocaudal basal nucleus following striatal injections in Cohort 2. Injection sites arranged from rostral (top) to caudal (bottom). All 6 cases resulted in many retrogradely labeled cells in the basal nucleus. Cell labeling in other nuclei (shaded in gray) is not shown for clarity. ***B.*** Chord diagram showing numbers of labeled cells in the Bpc, Bi, and Bpc by injection site placement. The top axis of this diagram shows the total number of labeled cells in Bpc (cyan), Bi (fuchsia), and Bmc (purple) across all cases examined. The bottom axis shows the number of labeled cells in each basal nucleus subdivision resulting from each striatal injection site, and is arranged counter-clockwise from the most rostral (left) to most caudal (right) injection site location. Each tick mark for both the top and bottom axes = 60 cells.

#### Direct cortical inputs to ‘limbic’ striatum regions

All 6 striatal cases contained labeled cells in both the PFC and insula (Fig. 8A). There were labeled cells in the agranular cortex after all injection sites. The injection site in the dorsomedial shell of the striatum resulted in labeled cells almost exclusively in the agranular cortices, while the injection site in the ventral striatal ‘core’ resulted in additional labeled cells in dysgranular cortices. Injections in more caudal sites of the ventral striatum led to more labeled cells in dysgranular and granular cortices. Overall, the greatest number of labeled cells from the PFC and insula were seen following injections into the rostral ventral striatum, with numbers tapering off following injections in more ‘caudal limbic striatum’.

**Figure 8:**
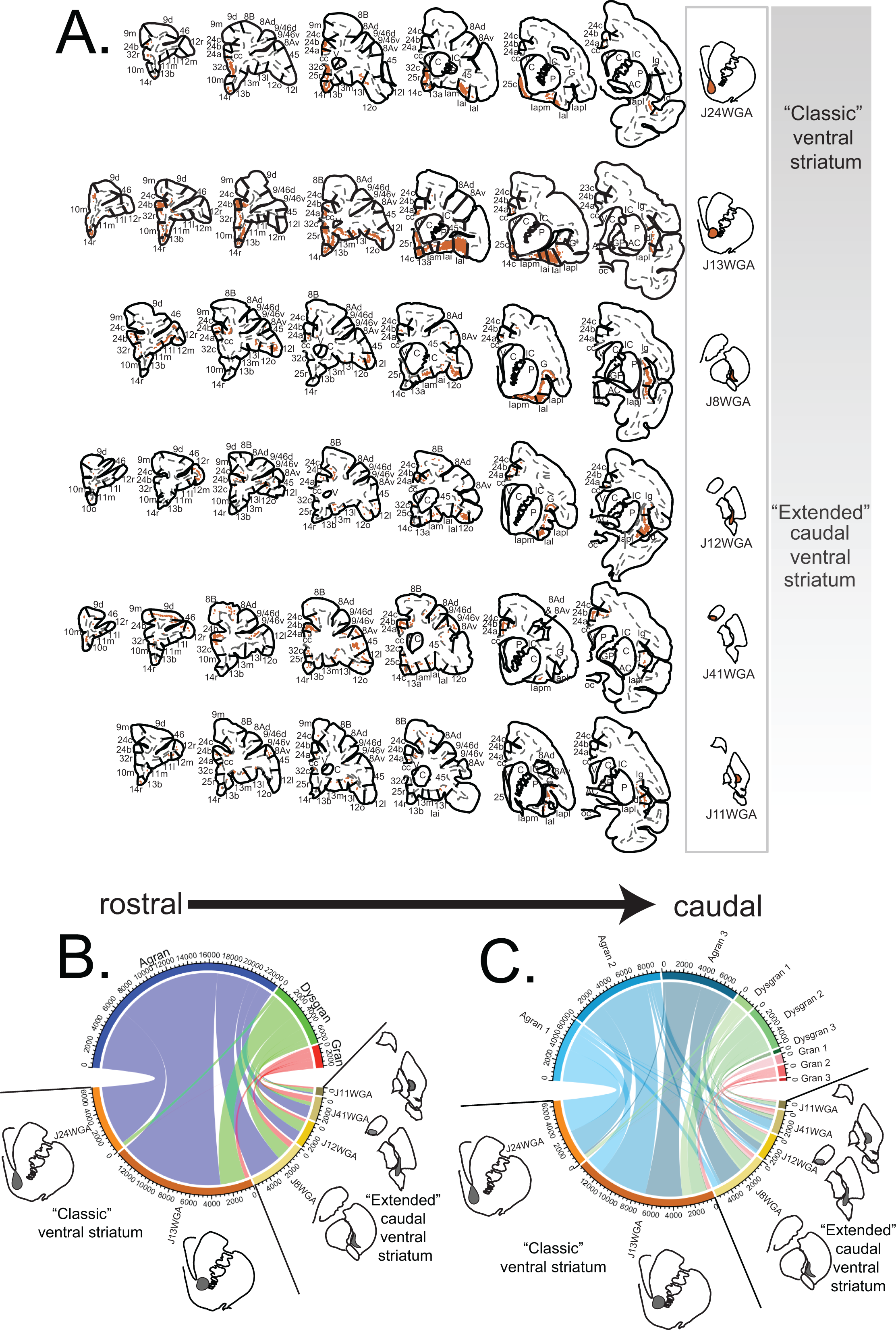
Cortico-’limbic’ striatal path. ***A.*** Representative charts depicting retrogradely labeled cells in the PFC and insula after injections in various striatal regions along the rostral-caudal extent of the ‘limbic’ striatum. Cases J24WGA and J13WGA had injections in shell and core of the ‘classic’ ventral striatum, respectively. Injection sites in J8WGA, J12WGA, J41WGA, and J11WGA were placed in different parts of the ‘extended’ ventral striatum. ***B.*** Chord diagram showing the number of retrogradely labeled cells (1:24 sections) in agranular (blue), dysgranular (green), and granular (red) cortical areas across all cases examined (top axis). The bottom axis shows the number of labeled cells found in agranular, dysgranular, and granular cortices after each striatal injection site. The bottom axis is arranged counter-clockwise from the most rostral (left) to the most caudal (right) injection site location. Each tick mark represents 400 cells. ***C***. Chord diagram using 9-group granularity analysis. following the same color schema in Figure 4C. Each tic mark for both the top and bottom axes represents 400 cells.

#### ‘Classic’ ventral striatum

Agranular cortices had many labeled cells in cases J24WGA and J13WGA, particularly areas 25c, 25r, 32c, 32r, and agranular insula (Fig. 8A). Both J24WGA and J13WGA had many labeled cells specifically in areas 25 and Iai, but J13WGA, placed in the ventral striatal core, had many additional labeled cells in Ial, as well as many labeled cells in agranular areas 14c and 24b, and dysgranular area 13b.

#### Caudal ventral striatum

Injections into the caudoventral putamen (J8WGA, J11WGA, and J12WGA) and the ventral body of the caudate head (J41WGA) resulted in labeled cells in the agranular cortices, as did our rostral ventral striatal site (Fig. 8A). The number of labeled cells from agranular cortices decreased along the rostrocaudal axis overall, balanced by increasingly more labeled cells appearing in the dysgranular and granular cortices. For example, cases J8WGA and J12WGA had a majority of cell labeling in agranular insula, but in cases J11WGA and J41WGA agranular insula labeling was more modest, with relatively more labeling in dysgranular insula. Similarly, the number of labeled neurons in agranular area 25 declined along the rostrocaudal axis. All caudal injection sites had many labeled cells in dysgranular insula, and moderate cell labeling in granular insula.

Quantitative analyses revealed hierarchical, nested projections based on basic levels of cortical differentiation (Fig. 8B). This pattern was similar to that observed in cortico-amygdala retrograde data, with agranular regions containing retrogradely labeled cells for all injection sites, and increasingly caudal regions containing labeled cells in dysgranular and granular cortices of both the insula and PFC. Analyses using a more refined (9 category) granularity classification showed similar patterns (Fig. 8C).

#### Comparing granularity index for cortico-amygdala-striatal and cortico-striatal Pathways

We used a ‘ratio of ratios’ approach to estimate how cortical granularity influenced amygdala projections to specific striatal regions in the ’indirect’ pathway (see Methods) and compared these results with results for the direct pathway (Fig. 9). For all striatal injection sites, agranular cortical sources dominated in both pathways, followed by a smaller contribution from dysgranular cortices, and the smallest from granular cortices (Fig. 9A). For the indirect pathway, input from each type of cortex remained relatively stable across the rostro-caudal extent of striatal injection sites. In contrast, in the direct pathway there was a shift in the relative contribution from agranular, dysgranular, and granular cortices along the rostrocaudal axis of the amygdala-recipient (’limbic’) striatum. In this path, the contribution from agranular cortical regions was progressively reduced, with increasing input from more differentiated cortices. (Fig. 9B & C). Consistent with these trends, there was relatively small variance in the percentage of labeled cells in agranular (29.602), dysgranular (9.975), and granular (5.403) cortex for the ’indirect’ path, in contrast to large variances in percentage of labeled cells in the agranular (627.216), dysgranular (300.639), and granular (127.727) cortices depending on the location of the injection site in the striatum (Fig. 9C).

**Figure 9:**
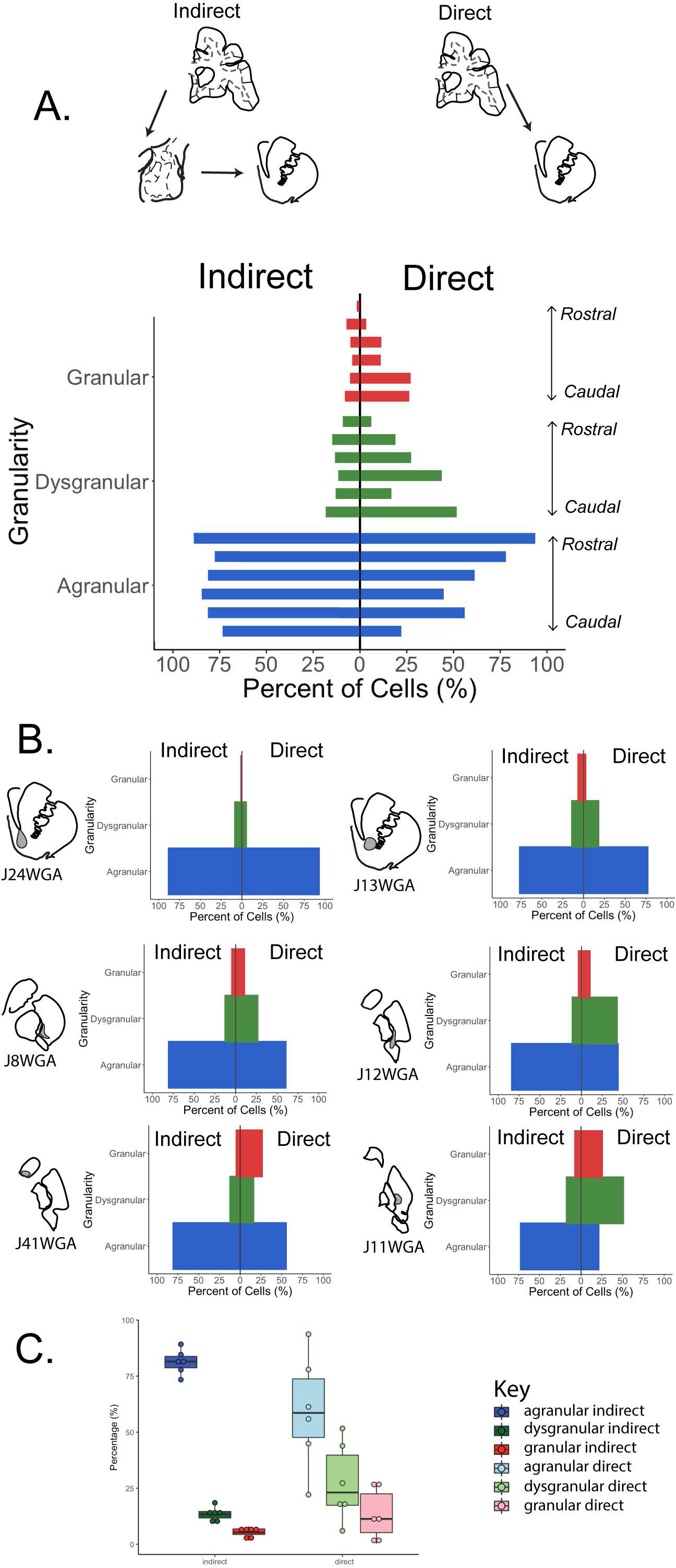
Indirect and direct pathway analysis**. *A.*** Composite diverging bar plots of each striatal injection showing the proportion of labeled cells in agranular (blue), dysgranular (green) and granular (red) cortices after calculating indirect pathway and direct pathway labeled cells. The proportion of labeled cells associated with amygdala inputs to all striatal sites (‘indirect’ path) is relatively consistent with a large percentage of cells categorized as agranular, and consistently small percentages of labeled cells categorized as dysgranular and granular, respectively. In contrast, the ‘direct’ cortico-‘limbic’ striatal path shows more variation with rostral ventral striatal sites having the greatest proportion of labeled cells in agranular cortices, and relatively small contributions from the dysgranular and granular cortices. This balance shifts at progressively caudal ventral levels, with reductions in labeled cells in agranular cortices, and incrementally more labeled cells in dysgranular cortices > granular cortices. ***B.*** Individual cases plotted for each path. ***C.*** Boxplots of agranular (blue), dysgranular (green) and granular (red) percentage data for indirect (dark shades) and direct (light shades) pathway data, illustrating differences in variance between granularity classes in the ’indirect’ and ’direct’ paths.

#### Cortical granularity maps: translating to traditional cortical divisions

After analyzing data using cortical granularity criteria, we were interested in understanding how the combinations of traditional cortical regions comprised these patterns. It has long been known that ’agranular’ cortices are associated with the ’limbic’ system, while dysgranular cortices are considered ’paralimbic’ and the most granular cortices are associated with higher cognitive functions (Badre and D’Esposito, 2009; Barbas, 2015). Since functional studies are typically based on cortical atlas designations, we ‘back-translated’ the granularity index findings by converting the data according into its original atlas designation (Table 2)(Fig. 10).

**Figure 10:**
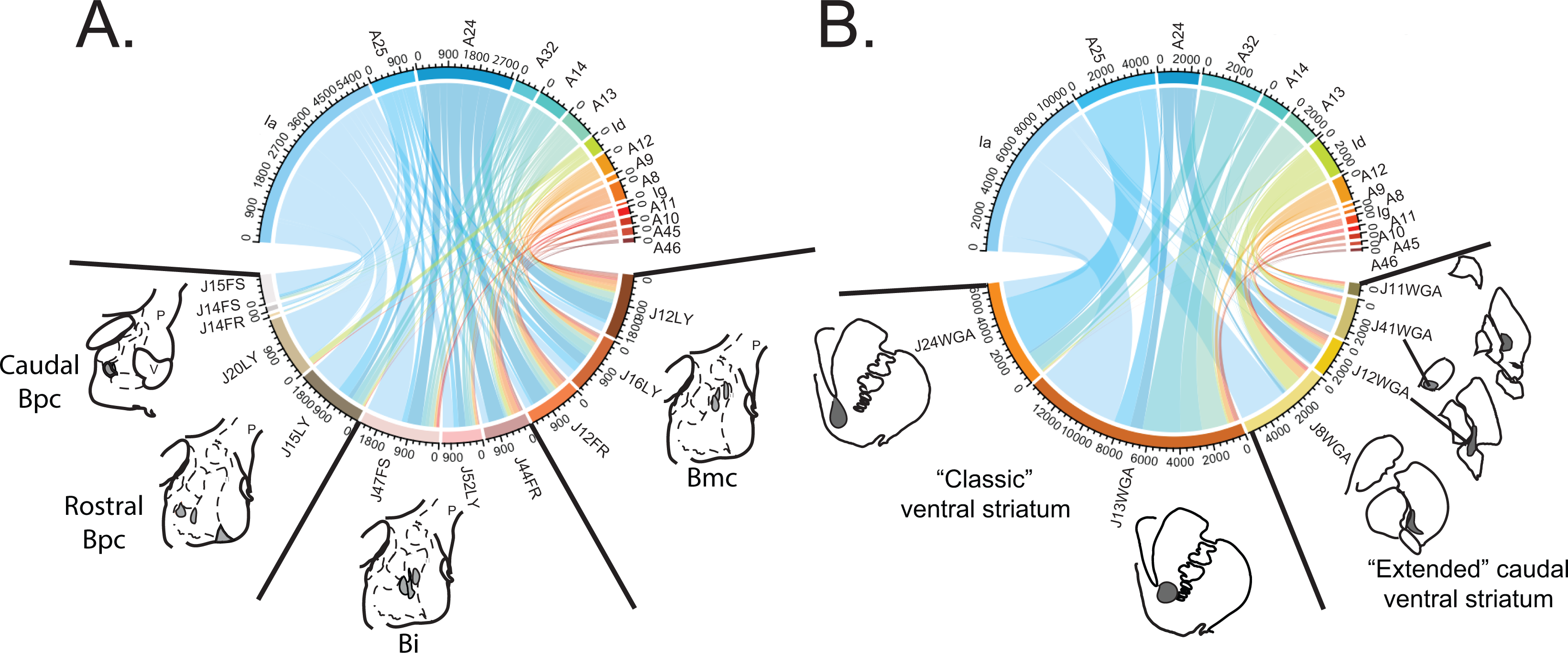
***A.*** Quantitative analysis of retrograde cortico-amygdala cell labeling in cortical areas (see text). The top axis shows the total number of labeled cells in each cortical area examined. Blue = agranular cortex only; blue-green = mix of agranular and dysgranular cortices, green = dysgranular cortex only; orange = mix of dysgranular and granular cortices, and red = granular cortex only. The top axis of this diagram shows the number of labeled cells from each cortical area resulting from each basal nucleus injection site. The bottom axis shows the number of labeled cells in each cortical region, per injection site. Each tic mark for both the top and bottom axes represents 180 cells. ***B.*** Quantitative analysis of retrograde cortico-striatal cell labeling by cortical area, indicated on the top axis, with the same color schema in ***A*.**. The bottom axis shows the number of labeled cells in each cortical region, resulting from each striatal injection site. Each tic mark for both the top and bottom axes represents 400 cells.

In the cortico-amygdala path, the greatest contribution to the ’agranular’ cortex was from agranular insula, where the numbers of labeled cells were greater than in all regions of the agranular cortex of the PFC combined (anterior cingulate areas 25, 24, and 32) (Fig 10A, blue). The agranular insula was associated with all basal nucleus subdivisions. In the Bpc, agranular insula was surprisingly dominant compared to area 25, especially in more caudomedial regions, based on cell count criteria. Labeled cells in area 25 resulted mainly from injection sites in the rostro-dorsal Bpc as well as the ventral Bi. In a similar manner, the number of labeled cells in areas 24 and 32 increased mainly from Bi and Bmc injection sites, having little association with Bpc injection sites.

Contributions from dysgranular cortices were mainly from dysgranular area 13, dysgranular area 8 (i.e. area 8B), dysgranular area 12 (i.e. 12o), and the dysgranular insula (Fig. 10A, aqua, green). Area 13 was associated mostly with Bmc injection sites, area 8 associated mostly with dorsal Bi and Bmc injection sites, area 12 associated mostly with dorsal Bi and Bmc injection sites, and dysgranular insula associated with rostral Bpc, dorsal Bi, and Bmc injection sites. In the granular cortices there were modest numbers of labeled cells in regions of the dorsal and lateral PFC (areas 9, 46, 45) and frontal pole (area 10). The granular insula (Ig) had the fewest labeled cells. All of these regions were associated mainly with the dorsal Bi and Bmc.

In the cortico-striatal path, the agranular cortex as a whole had its largest contribution from the agranular insula, followed by area 25. Labeled cells in the agranular insula were associated with all striatal sites, while labeled cells in area 25 were associated mainly with rostral ventral striatal injection sites. Areas 24 and 32 contained labeled cells in all striatal injection sites, which were prominent in the rostral striatum, but persisted in association with every caudal ventral striatal site. Contributions from the dysgranular component of area 13 (i.e. area 13b) (aqua) and dysgranular insula (light green) were first seen in the core of the rostral ventral striatum, and were present through the caudal ventral striatum. The dysgranular insula (light green) was a key contributor following all caudal ventral striatal injection sites. The number of labeled cells in combined dysgranular/granular (orange/red) cortices was most prominent in area 12 (specifically area 12o), with lesser contributions from other combined dysgranular/granular and fully granular cortices. These findings are consistent with previous work in monkey (Selemon and Goldman-Rakic, 1985; Yeterian and Pandya, 1991; Choi et al., 2017a; Choi et al., 2017b). Labeled cells in all these regions were associated with the core of the rostro-ventral striatum, and caudal ventral striatal sites. The ’core’ of the rostral ventral striatum had the most labeled cells overall, compared to other injection sites. Labeled cells from the PFC and insula generally declined in ’extended’ caudal ventral striatal regions. These regions receive additional massive inputs from sensory association cortices not examined in this study (Saint-Cyr et al., 1990; Yeterian and Pandya, 1995, 1998) (see Discussion).

#### Cortico-amygdala k-means cluster analysis

We then took a machine learning approach to examine how granularity-specific cortical subregions project to discrete regions of the basal nucleus and striatum (Fig. 11A-F). For cortico-amygdala retrograde data, we found differences in how cortical regions cluster across the caudal-ventral and rostral-dorsal extent of the basal nucleus, and were able to further examine cluster identities by inspecting tables containing the z-scores of all cortical regions per cluster (Fig. 11A-C, Fig. 11-1.) Four clusters were selected, based on criteria for silhouette analysis. Three different clusters were composed of distinct agranular cortices, and one cluster comprised of multiple agranular, dysgranular, and granular cortices (Fig. 11B & C). The ‘Agranular mosaic’ cluster (Fig. 11B, green) is an exclusively agranular cluster that contains areas Iapm, 25c, Iai, and Ial. Large z-scores characterized all cortical regions in this cluster (reflecting many retrogradely labeled cells greater than the mean), all of which are associated with all basal nucleus subdivisions. There was a preponderance of ‘high’ and ‘mean/above mean’ z-scores for retrogradely labeled cells in cortical areas associated with the Bpc and Bi, and some ‘mean/above mean’ z-scores existing in the Bmc.

**Figure 11:**
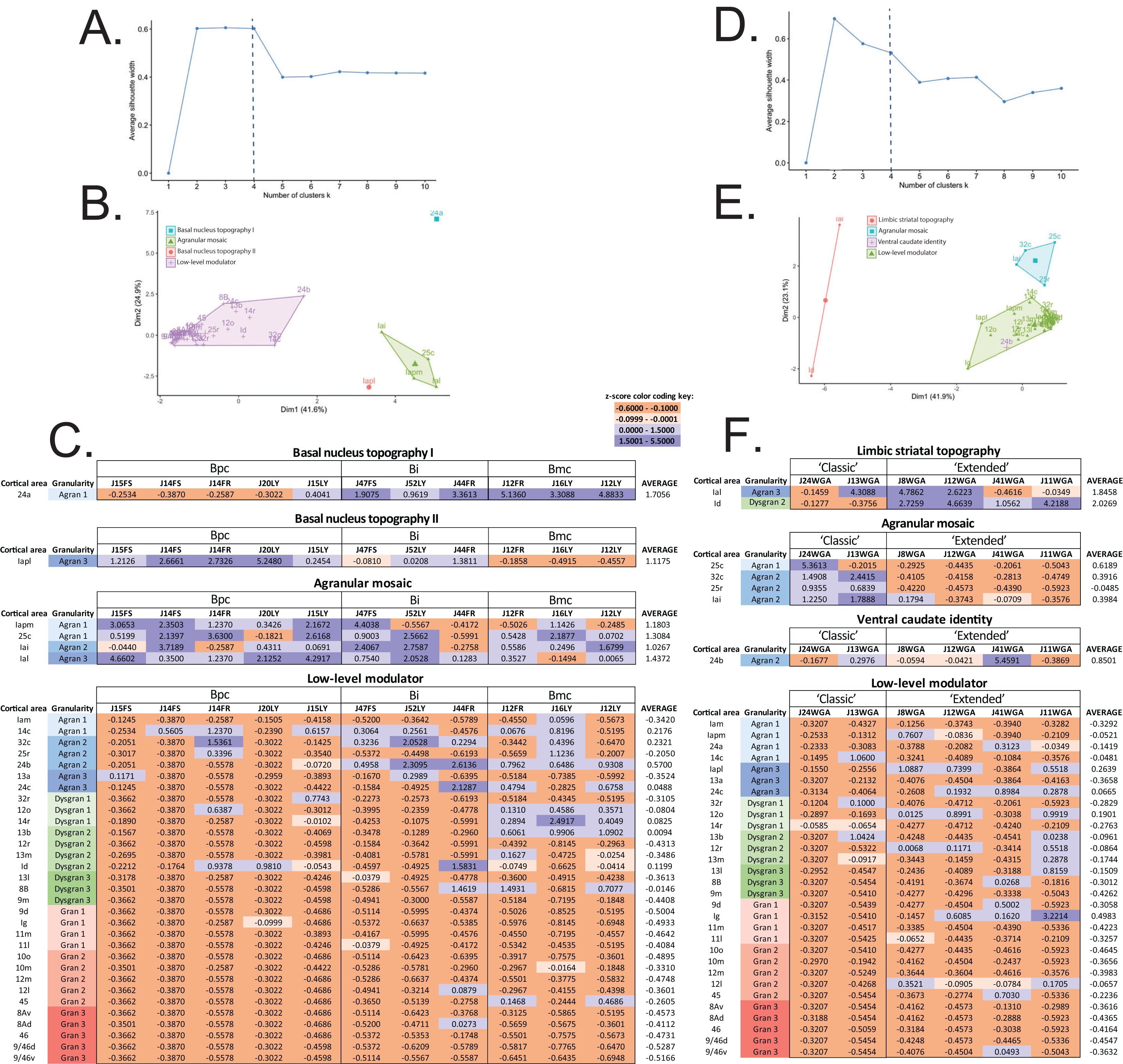
K-means cluster analyses for cortico-amygdala (A-C) and cortico-striatal (D-F) data. ***A.*** Silhouette plot for cortico-amygdala data. ***B*.** K-means cluster analysis plot for cortico-amygdala data. Each axis represents the percentage of the data captured in the two dimensions that account for the highest variance across the data. 4 clusters are named by descriptive labels, based on which feature of the data they capture: ‘Basal nucleus topography I’ (teal), ‘Agranular mosaic’ (green), ‘Basal nucleus topography II’ (red-orange), and ‘Low-level modulator’ (lavender). ***C.*** Cortico-amygdala cluster identities, based on inspection of z-scores across each cluster. Table columns are organized from caudal-ventral to rostral-dorsal injection site location. Z-scores are grouped from ‘high’ to ‘low’ values: ‘high’ = mean z-score between 1.5001 – 5.5000 (dark purple), ‘mean/above mean’ = mean z-score between 0.0000 – 1.5000 (light purple), ‘slightly below mean’ = mean z-score between -0.0999 - -0.0001 (light orange), and ‘largely below mean’ = mean z-score between -0.6000 - -0.1000 (dark orange). ***D.*** Silhouette plot for cortico-striatal data. ***E.*** K-means cluster analysis plot for cortico-striatal data, with 4 clusters identified by descriptive labels: ‘Limbic striatal topography’ (red-orange), ‘Agranular mosaic’ (teal), ‘Ventral caudate identity’ (lavender), and ‘Low-level modulator’ (green). ***F.*** Cortico-striatal cluster identities, based on inspection of the distribution of z-scores for each cluster. Extended data Figure 11-1 shows a table listing the mean and standard deviation for cell counts for each injection site for calculating z-scores.

**Figure 11-1:**
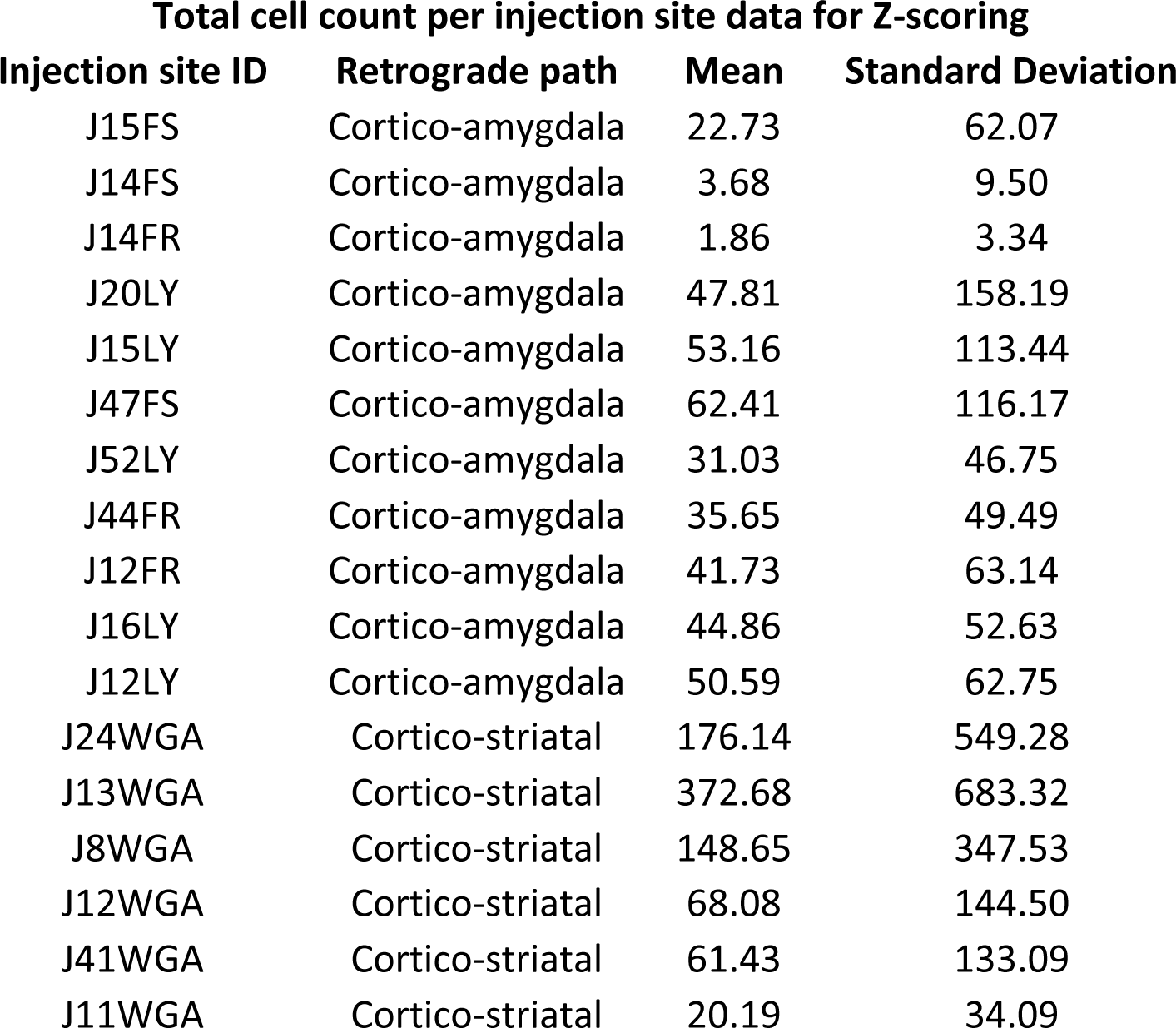
Means and standard deviations of total labeled cortical cell counts found for each basal nucleus and striatal injection site.

There were two small clusters best defined as ’Basal nucleus topography’ clusters (Fig. 11B, blue and red). One consisted exclusively of agranular area 24a (’Basal nucleus topography I’, teal) and identified the relative dorsal position of the injection in the basal nucleus. The ’Basal nucleus topography I’ cluster had labeled cells with increasingly high z-scores ranging from ’largely below mean’ z-scores in Bpc sites, to values of ‘high’ z-scores and ‘mean/above mean’ z-scores (i.e. z-scores between 0.0000 - 1.5000) for Bi sites, and ‘high’ z-scores for Bmc sites and the Bi site (J44FR) nearest the Bmc (i.e. z-scores between 1.5001 - 5.5000) (Fig. 11C). In contrast, the ’Basal nucleus topography II’ cluster (Fig. 11B, red-orange) was exclusively composed of agranular cortical region Iapl, and was associated with a basal nucleus injection location in the opposite direction, from ventral to dorsal. This small cluster reflected retrogradely labeled cells with ‘high’ and ‘mean/above mean’ z-scores for Bpc sites, ‘mean/above mean’ z-scores and ‘slightly below mean’ z-scores (i.e. z-scores between 0.0999 - -0.0001) for Bi sites, and ‘largely below mean’ z-scores for Bmc sites. Together, ’Basal nucleus topography clusters I and II’ are inversely related, and gradually increase in their contributions along the ventral-dorsal and dorsal-ventral extent of the basal nucleus, respectively (Fig. 11C).

The ‘Low-level modulator’ cluster (Fig. 11B, lavender) comprised many agranular, dysgranular, and granular cortical areas which had ‘largely below mean’ z-scores across all basal nucleus injection sites. This cluster describes multiple, low to modest, contributions from labeled neurons in specific cortical projections, the combinations of which define specific basal nucleus subdivisions (Fig. 11C). While all basal nucleus subdivisions received multiple ’low level’ inputs, the dysgranular and granular cortices had relatively greater numbers of labeled cells (reflected by z-scores above the mean) associated with the Bi and Bmc compared to the Bpc.

#### Cortico-striatal k-means cluster analysi

(Fig. 11D-F, Fig. 11-1). Using the silhouette method as a guide for selecting an optimal *k*, cortical connectivity with the striatum was best represented through four clusters (Fig. 11D). Similar to cortico-amygdala data, clusters are described after analyzing the z-scores of retrogradely labeled cells in all cortical regions within a cluster, and their relationship to connections with the striatum (Fig. 11E & F). In this dataset, there were two agranular clusters, one agranular/dysgranular cluster, and one cluster containing agranular, dysgranular, and granular inputs (Fig. 11E & F).

The ‘Agranular mosaic’ cluster consisted of agranular cortical areas Iai, 32c, 25c, and 25r (Fig. 11E, teal). This cluster revealed key differences between the ‘classic’ rostral ventral striatum (revealing retrogradely labeled cells in these cortical regions with ‘high’ and ‘mean/above mean’ z-scores) and the ‘extended’ caudal ventral striatum, (where the numbers of retrogradely labeled cells were relatively lower, reflected in z-scores ‘largely below mean’) (Fig. 11F). In the classic ventral striatum shell (J24WGA), area 25c was a strong, defining input, revealed by an exceptionally ‘high’ z-score for retrogradely labeled neurons, compared to all other regions. Notably, area 25c had weaker input to the core (J13WGA). In sum, the ’Agranular mosaic’ cluster differentiated the ‘classic’ ventral striatum from the ‘extended’ caudal ventral striatum.

Inspection of z-scores for a cluster containing the Ial (agranular) and Id (dysgranular), revealed a topographic distribution, leading to its description as a ‘Limbic striatal topography’ cluster (Fig. 11E, red-orange). Ial projections had ‘high’ z-scores (i.e. z-scores between 1.5001 - 5.5000) in the classic ventral striatum ‘shell’, which dropped progressively across more caudal ventral striatal sites (Fig 11F). The opposite relationship occurred for Id, where projections associated with caudal ‘extended’ striatal sites (J8WGA, J12WGA, and J11WGA) have ‘high’ z-scores, and progressively drop as injections moved to the ventral caudate (J41WGA) and to classic ventral striatal sites. This inverse rostral-caudal shift from ‘high-low’ Ial influence to ‘low-high’ Id influence occurs posterior to the anterior commissure in the ventral putamen (J8WGA and J12WGA).

One small cluster consisting of agranular area 24b, was uniquely associated with the injections in the body of the caudate nucleus; we termed it the ‘Ventral caudate identity’ cluster (Fig 11E, lavender). This cortical region had an exceptionally ‘high’ z-score in case J41WGA (5.4591), which was substantially lower for all remaining sites (Fig. 11F). This strongly suggests that area 24b is a defining input to the ventral body of the caudate nucleus, relative to other ’limbic striatal’ sites.

Similar to the cortico-amygdala path, there was a ‘Low-level modulator’ cluster for the cortico-striatal path (Fig. 11E, green, 11F). Similar to the cortico-amygdala path, z-scores of retrogradely labeled cells showed relatively modest to weak contribution across agranular, dysgranular and granular cortices for all striatal sites. Also similar to the cortico-amygdala path, while all striatal subdivisions received multiple ’low level’ inputs, the dysgranular and granular cortices inputs were shifted, and were associated with relatively greater numbers of labeled cells (reflected by z scores above the mean, purple) associated with the ’extended’ caudal ventral striatum compared to the ’classic’ ventral striatum.

## Discussion

Several important findings emerged from these studies. The first is that a ‘cortical logic’ governed by laminar structure shapes information flow in both the ‘indirect’ and ‘direct’ paths to the ’limbic’ striatum (Fig. 12). Agranular cortical inputs to the basal nucleus of the amygdala and striatum were foundational, as evidenced by the presence of agranular cortical inputs throughout both the cortico-amygdala and cortico-striatal paths. These agranular cortical areas undergird inputs, first from more dysgranular cortex, and then from more granular cortices. In addition, while not a laminar structure, the basal nucleus is cortical-like (Carlsen and Heimer, 1988), and appears to follow hierarchical rules found in cortical projections to the striatum.

**Figure 12:**
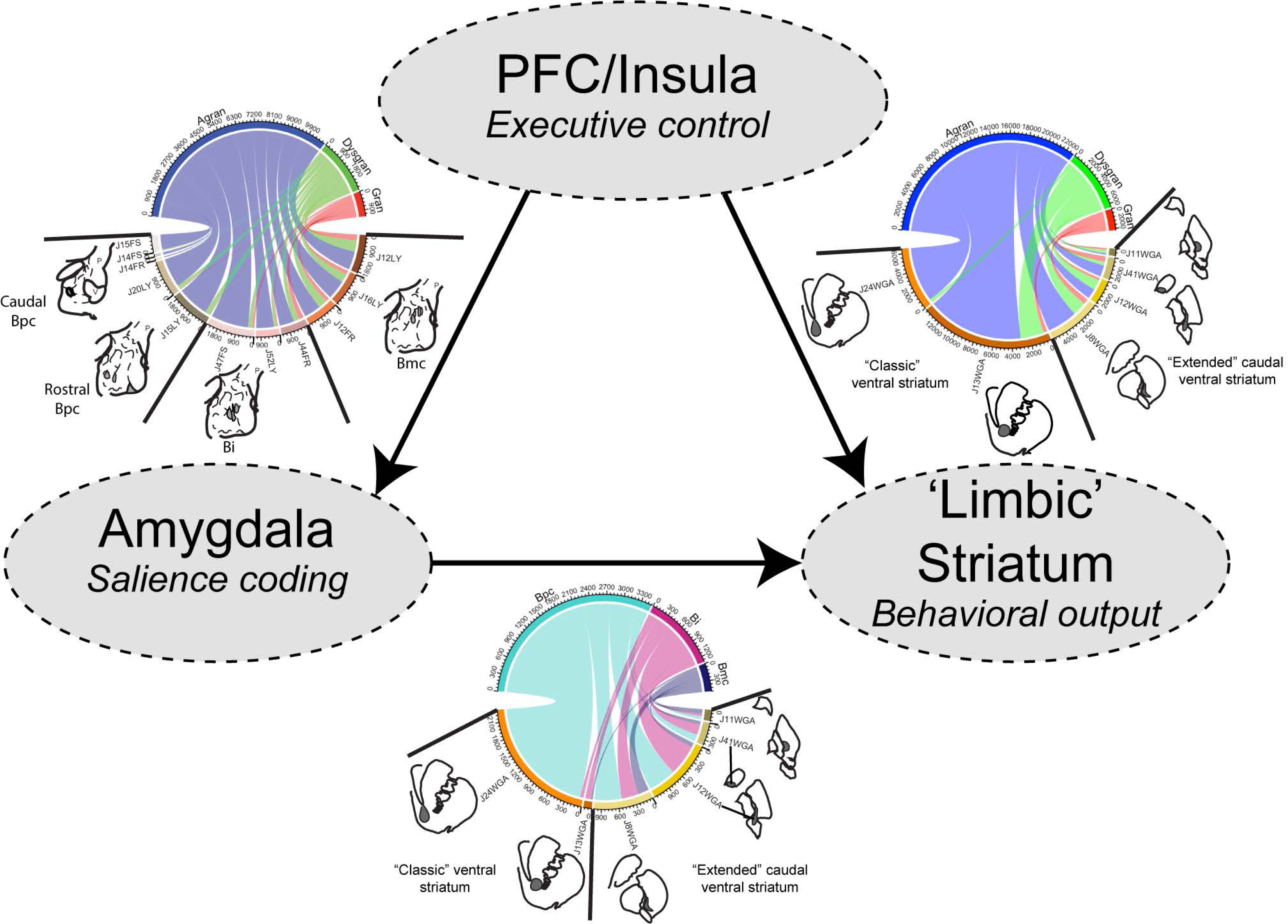
Overview of ‘cortical logic’ in the ‘indirect’ pathway (PFC/Insula → amygdala connectivity, left + amygdala → ‘limbic’ striatum connectivity, bottom) and the ‘direct’ pathway (PFC/Insula → ‘limbic’ striatum connectivity, right).

Another key result was that cortico-amygdala-striatal paths were generally influenced by a consistent proportion of agranular/dysgranular/granular cortices, overwhelmingly dominated by agranular cortex, regardless of whether the final target was the rostral-ventral or caudal-ventral striatum. In contrast, the ratio of agranular/dysgranular/granular cortical input in the direct cortico-amygdala projection was more variable, and gradually tipped to favor fewer agranular, and more differentiated inputs in the caudal ventral striatum. These results were supported using machine learning approaches in cluster analyses.

Finally, we concluded that the ‘limbic’ striatum, as defined by amygdala inputs, extends beyond the ’classic’ ventral striatum (nucleus accumbens), as has been previously noted (Russchen et al., 1985; Fudge et al., 2004; Choi et al., 2017a). Different rostrocaudal levels of the ‘limbic’ striatum receive shifting combinations of cortical inputs, making for unique ‘connectional fingerprints’ in each region.

### Cortical logic

Many individual studies have examined cortico-amygdala, amygdalo-striatal, and cortical-striatal pathways, using anterograde tracer injections (Mufson et al., 1981; Russchen et al., 1985; Goldman-Rakic and Selemon, 1986; Carmichael and Price, 1995; Ferry et al., 2000; Ghashghaei and Barbas, 2002; Fudge et al., 2004). Our work builds on these studies by taking a ‘connectomics’-type approach, examining the relationship of multiple pathways within and across animals, and assessing rules by which top-down cortex modulates down-stream targets. This approach is based on a basic tenet that functional specificity is determined by ensembles of afferent projections, and that the strength of any one connection depends on features of the afferents that arrive with it.

The principles that guide the ’logic’ of cortical afferents are based in the granular complexity of the cortical afferent source. The least differentiated cortical regions examined form a ‘foundational’ base across the basal nucleus and ‘limbic’ (amygdala-recipient) striatum, while inputs from more differentiated cortical regions are additive and are always seen as co-projections with these less differentiated cortical regions. Cluster analyses for both the cortico-amygdala and cortico-striatal paths support this overarching principle, and also highlighted subtle differences between the pathways. For example, in the cortico-amygdala path, an ’agranular mosaic’ cluster revealed strong inputs to all of the basal nucleus. In contrast, in the cortico-striatal path, while agranular cortex projected throughout the striatum, two clusters divided it. The ‘agranular mosaic’ was high only in the ’classic ventral striatum’ while the ‘Limbic striatal topography’, consisting of agranular and dysgranular inputs, predicted the location of injection sites along the rostro-caudal axis. In accord with general ’cortical logic’, an overlay of multiple dysgranular and granular inputs (‘low-level modulator’ clusters) was a specific feature of each pathway, added only in relatively dorsal (Bmc) or caudal (‘extended’ ventral striatum) sectors.

### “Connectivity Fingerprints” and functional anatomic implications

Due to the ‘logic’ of cortical afferents, a predictable, yet unique set of inputs are found in specific subregions of both the basal nucleus and the striatum. Unique connectivity profiles (’fingerprints’) are found in ventral-dorsal (basal nucleus) and rostral-caudal (striatum) locations, and shift in a predictable, topographical manner.

### Basal nucleus ‘fingerprints

Against a backdrop of broad-based agranular cortical inputs, projections from increasingly dysgranular and granular cortical regions are added, and overlapped in incrementally dorsal and rostral aspects of the basal nucleus. The ’agranular mosaic’ cluster common to all basal nucleus regions (Iapm, 25c, Iai, Ial) suggests that interoceptive information and autonomic functions modulate the basal nucleus as a whole (Craig, 2002; Wallis et al., 2017). Interestingly, agranular areas 24a and Iapl increase and decrease, respectively, along the Bpc-Bi-Bmc axes, providing a connectional marker of this gradient (Basal nucleus topography I and II clusters). Areas 24b, 12o, 13b, 8B, 45 (found in the ’low level modulator’ cluster) contribute increasingly dysgranular and granular cortical inputs in the Bi and Bmc; these connections are implicated in updating the value of rewards during decision-making (Murray and Rudebeck, 2018).

#### ‘Limbic’ striatal ‘fingerprints’

The general connectional rules found are also obeyed in the ’limbic’ striatum, but with sharper shifts in cortical inputs, suggesting clear functional changes along the rostrocaudal axis. The classic ventral striatum (also known as the nucleus accumbens) is well-known to mediate reward-driven behaviors (Haber and Knutson, 2010) based in part on amygdala inputs (Popescu et al., 2007; Dallerac et al., 2017). The ‘classic’ ventral striatum also received massive inputs from 25c, 32c, 25r, Iai (captured in the ’agranular mosaic’ cluster), which tapered sharply in the ‘extended’ ventral striatal sites. Within this agranular cluster, the dorsomedial shell of the nucleus accumbens has a restricted, high input from area 25c, consistent with previous work (Kunishio and Haber, 1994; Chikama et al., 1997). Together, these data inputs from a tight, interconnected cortical network associated with interoceptive processing (Carmichael and Price, 1996) that is important for reward-driven responses.

Agranular cortical inputs found in the classic ventral striatum persist through the caudal ventral (’extended’) striatum but decline, as dysgranular and granular cortical inputs are increasingly added. This pattern is captured in the ‘Limbic striatal topography’ cluster, which shows an increasing input from Id (important for social responses (Jezzini et al., 2012)), review (Evrard, 2019)) with a decreasing input from Iapl, demarcating a gradual connectional boundary between the rostral and caudal ventral striatum. Notably, basal amygdala inputs to the striatum follow similar cortical rules. Bpc projects broadly throughout the ’limbic’ striatum, maintaining inputs to the caudal ventral striatum, which are reduced as additional inputs from Bi, and particularly, Bmc enter. Bmc, which itself receives the most differentiated cortical inputs from the orbital and ventrolateral prefrontal cortex (Fig. 12), exclusively projects to the caudal ventral striatum, a recipient of these direct cortical inputs (i.e. areas 24c, 12l, 12o, 45, Id/Ig) (Selemon and Goldman-Rakic, 1985; Saint-Cyr et al., 1990; Yeterian and Pandya, 1991; Fudge et al., 2005; Amita et al., 2019). The caudal ventral striatum has long been known as the ’sensory’ striatum due to prominent inputs from auditory and visual association cortices in the temporal lobe (Yeterian and Pandya, 1995, 1998). Here we show that these caudal ventral striatal areas also receive inputs from specific cortical areas involved in awareness of bodily movements (Ig, 24c) (Morecraft and Van Hoesen, 1992; Karnath and Baier, 2010; Jezzini et al., 2012) see review (Evrard, 2019) object identification (area 12) (Wilson et al., 1993), and social communication (areas 12l, 45) (Romanski, 2012) via ’indirect’ and ’direct’ paths. Taken together with established caudal-ventral striatal visual/auditory inputs in primate species, this connectional fingerprint suggests a role in motivated multisensory responses, including complex social interactions.

#### Driver/modulators

One way to view the broad ‘foundational’ influence of agranular cortices versus the relatively more restricted influence of differentiated cortices is by applying emerging ideas about differential functions of cortical afferent systems (Sherman and Guillery, 1998). This concept is based on Sherman and Guillery’s ‘driver-modulator’ theory of excitatory afferents, in which a ‘driver pathway’ provides the main path for information flow and a co-projecting ‘modulator pathway’ regulates the output from ‘driver’ systems (Sherman and Guillery, 1998). Applied to our work here, we hypothesize that the agranular cortices comprise driver systems in ’indirect’ cortico-amygdala-striatal and ’direct’ paths due their widespread presence in these target structures. In the ’indirect path’, the basal nucleus—itself a ’cortical-like’ structure without lamination—projects strongly throughout the ’classic’ and ’caudal ventral’ striatum, and may be an important ’driver’. The other cortical inputs composed of varying combinations of dysgranular and granular cortices point to specific micro-targets along the ventral-dorsal and rostro-caudal axis in both the amygdala and striatum, respectively. These most differentiated cortical regions may be ‘modulator’ pathways, based on their generally lighter inputs, and the fact that they are always present with denser inputs from less differentiated cortical areas. Specific anatomic and electrophysiologic properties that define ’driver/modulator’ paths can be further probed in these systems at the physiologic and anatomic levels (Sherman and Guillery, 1996; Lee and Sherman, 2009; Kelly et al., In Press).

#### Conclusion

Cortical granularity rules were elucidated in a well-known triad of connections through the ‘limbic’ brain in monkeys. Using a connectomics-type approach, we found that the basal nucleus and ‘limbic’ striatum are not homogeneous entities, and have unique, predictable, sets of ‘connectivity fingerprints’ within them. These general connectivity patterns have implications for the ways in which the emotional brain can code increasingly complex levels of social and cognitive information in a flexible manner, such as predicting actions and choices of others (Saez et al., 2015; Grabenhorst et al., 2019), and may help us to better understand dynamics in complex neuroanatomic circuits associated with human psychiatric disease.

## Acknowledgements

This work was funded through the support of the National Institutes of Mental Health R01MH63291 (J.L.F.), and the Schmitt Program on Integrative Neuroscience (SPIN) GR504304 (J.L.F.). We thank Nanette Alcock for assistance with histology and immunocytochemistry. We also thank Keshov Sharma for R code advice and helpful discussions about data analysis.

